# β-hydroxybutyrate is a metabolic regulator of proteostasis in the aged and Alzheimer disease brain

**DOI:** 10.1101/2023.07.03.547547

**Authors:** SS Madhavan, S Roa Diaz, S Peralta, M Nomura, CD King, A Lin, D Bhaumik, S Shah, T Blade, W Gray, M Chamoli, B Eap, O Panda, D Diaz, TY Garcia, BJ Stubbs, GJ Lithgow, B Schilling, E Verdin, AR Chaudhuri, JC Newman

## Abstract

Loss of proteostasis is a hallmark of aging and Alzheimer disease (AD). Here, we identify β-hydroxybutyrate (βHB), a ketone body, as a regulator of protein solubility in the aging brain. βHB is a small molecule metabolite which primarily provides an oxidative substrate for ATP during hypoglycemic conditions, and also regulates other cellular processes through covalent and noncovalent protein interactions. We demonstrate βHB-induced protein insolubility across *in vitro*, *ex vivo*, and *in vivo* mouse systems. This activity is shared by select structurally similar metabolites, is not dependent on covalent protein modification, pH, or solute load, and is observable in mouse brain *in vivo* after delivery of a ketone ester. Furthermore, this phenotype is selective for pathological proteins such as amyloid-β, and exogenous βHB ameliorates pathology in nematode models of amyloid-β aggregation toxicity. We have generated a comprehensive atlas of the βHB-induced protein insolublome *ex vivo* and *in vivo* using mass spectrometry proteomics, and have identified common protein domains within βHB target sequences. Finally, we show enrichment of neurodegeneration-related proteins among βHB targets and the clearance of these targets from mouse brain, likely via βHB-induced autophagy. Overall, these data indicate a new metabolically regulated mechanism of proteostasis relevant to aging and AD.

## INTRODUCTION

Alzheimer disease (AD) remains one of the most impactful human diseases with few effective disease-modifying therapies. AD is characterized by deficits in brain energy metabolism and altered protein homeostasis (proteostasis)^1–5^. These perturbations in metabolism and proteostasis are present in sporadic cases and further exacerbated by genetic regulators of the disease process, such as APOE alleles^6–10^. The multifactorial nature of AD contributes to the difficulty in developing effective therapies, as the connections between metabolism and proteostasis are poorly understood. Moreover, the dominant risk factor for developing AD is chronological age^3^. Aging is also characterized by loss of proteostasis and deregulated nutrient-signaling, among a number of other molecular mechanisms that are known to be connected metabolically^11–13^. Small molecule metabolites can link metabolism with aging mechanisms through secondary protein-interacting roles, in addition to their core energetic functions. Here, we investigate direct proteostatic effects of a cellular metabolic state induced via small molecule metabolites.

Ketone bodies are a class of hepatically-sourced and lipid-derived small molecule metabolites which include acetone, acetoacetate, and (R)-β-hydroxybutyrate (R-βHB)^14–17^. The primary function of acetoacetate and R-βHB production is to provide cellular energy in extrahepatic tissues during periods of hypoglycemia, such as fasting, starvation, high-intensity exercise, and ketogenic diet. Ketone bodies can also be administered exogenously without nutritional changes, such as via a ketone ester. Beyond providing energetic substrate, R-βHB also possesses direct covalent and non-covalent protein-binding activities, including inhibition of histone deacetylases, post-translational modification of histone and non-histone proteins, inhibition of the NLRP3 inflammasome, and binding to cell surface receptors^14, 18–26^.

There is clear preclinical literature support, and early clinical data, for ketogenic therapies in aging and AD^27^. A non-obesogenic ketogenic diet extends lifespan in mouse models and improves healthspan outcomes, including memory, in aged mice^28–30^. Additionally, ketogenic diet and exogenous ketones have been shown to improve cognitive and motor behavior in several mouse models of AD^31–34^. Early human studies of ketogenic compounds have improved cognitive scores in patients with mild to moderate AD^35–40^. While the molecular mechanisms underlying these improvements in the aging and AD brain are not fully clear, and may be multifactorial, there has been preliminary evidence for an effect of ketone bodies on proteostasis. R-βHB directly prevents amyloid-β_1-42_ toxicity in rat hippocampal cells, and both ketogenic diet and exogenous ketones reduce total plaque burden in the brain of multiple mouse models of AD^31, 41–43^. Reduction of total plaque burden in the brain of mouse models of AD has also been replicated in other ketosis-inducing dietary interventions, such as calorie restriction and intermittent fasting^44, 45^. Multiple recent studies have implicated the role of ketone bodies in regulating autophagic flux and chaperone-mediated autophagy, including clearance of amyloid-β and pathogenic tau^46–50^. However, a clear mechanism by which βHB directly interacts with proteins or protein clearance machineries has not been identified.

The deposition of misfolded proteinaceous aggregates resulting from a loss in proteostasis is a hallmark of aging and neurodegenerative diseases (NDDs), including AD^1, 3, 5, 11–13^. The relative solubility of these misfolded proteins within the brain, especially in AD, is of clinical importance, as soluble oligomeric proteins exhibit prion-like properties^1, 5^. These soluble oligomers seed aggregation and spread from cell-to-cell throughout the brain as a marker of disease progression^1, 51, 52^. Indeed, although the role of soluble oligomers versus protein aggregates in AD pathophysiology is unsettled, the insolubilization of these misfolded oligomeric proteins, particularly if chaperoned to degradation, may act as a barrier to the progression of these AD pathology and may be a mechanism of cellular damage control in NDDs. Notably, soluble oligomer-targeted antibody therapies such as lecanemab have recently shown clinical success^53, 54^.

Here, for the first time, we identify a novel direct protein-interacting function of βHB and structurally similar small molecule metabolites in proteostasis. We report selectivity in this proteostatic regulation for pathogenic proteins such as amyloid-β_1-42_, and evidence of amelioration of amyloid-β_1-42_ toxicity *in vitro* in mammalian cells and *in vivo* with multiple *C. elegans* strains. Furthermore, we generated libraries of protein targets both *ex vivo* and *in vivo* from aged mouse brain via data-independent acquisition mass spectrometry. We observed enrichment for NDD-related proteins in both *ex vivo* and *in vivo* libraries. Finally, we show that βHB-induced insolubility leads to misfolded protein turnover *in vivo*, likely via βHB communication with cellular protein degradation pathways. This work identifies βHB as a global regulator of cytosolic protein solubility, and identifies new metabolism-related mechanistic targets for therapeutic development in aging and AD.

## RESULTS

### β-hydroxybutyrate directly induces protein insolubility without posttranslational modification

We, and other laboratories, have shown that memory phenotypes in aged mice and mouse models of Alzheimer disease (AD) can be improved with ketogenic therapies^28–32^. While the substrate provision for brain energy metabolism is likely indirectly assisting with these improvements, no direct mechanism has been validated. As previous literature has identified a clear connection between ketosis and clearance of AD plaque burden in mouse models of AD^31, 42, 43^, and ketone bodies are known posttranslational modifiers^18–22^, we became interested in the potential direct effects of β-hydroxybutyrate (βHB) on misfolded proteins.

Given the well-known effects of pH on protein solubility, and since βHB and many other small molecule metabolites are organic acids, we used a working buffer (TEM), that preserved physiological pH despite addition of acidic metabolite compounds (Extended Data 1a-b). Additionally, we carefully pre-buffered protein solutions and lysates prior to the addition of compounds to prevent local changes in pH (i.e. first pipetting metabolite, then TEM buffer, then TEM-buffered protein solutions or lysates).

We used centrifugation to pellet proteins whose solubility in TEM had changed after incubation, then resolubilized this protein pellet with a mixed detergent buffer (NDSD) prior to further analyses. Full details of buffers are covered in Materials and Methods, and all chemical structures of compounds tested are shown in Extended Data 1c.

We began in a highly purified system with bovine serum albumin (BSA) as a test protein, whose misfolding can be predictably induced with heat^56^. We co-incubated BSA with and without 10 mM R-βHB and a structurally similar ketogenic alcohol 1,3- butanediol (1,3-BD) at +37°C (native folding) and +70°C (heat-induced misfolding), then used centrifugation to separate soluble and induced-insoluble fractions before resolubilizing insolubilized proteins in NDSD buffer (Fig. 1a). Using bis-ANS fluorescence, which increases with exposure of non-polar cavities in proteins^57^, we confirmed heat-induced misfolding of BSA at +70°C (Extended Data 2a). We found that most native folded BSA remained soluble, regardless of compound addition, as did heat-misfolded BSA without compound addition. However, R-βHB induced insolubility of heat-misfolded BSA by approximately 3-fold, while the structurally similar alcohol analogue 1,3-BD did not, as quantified by Imperial staining (Fig. 1b).

**Fig. 1.**
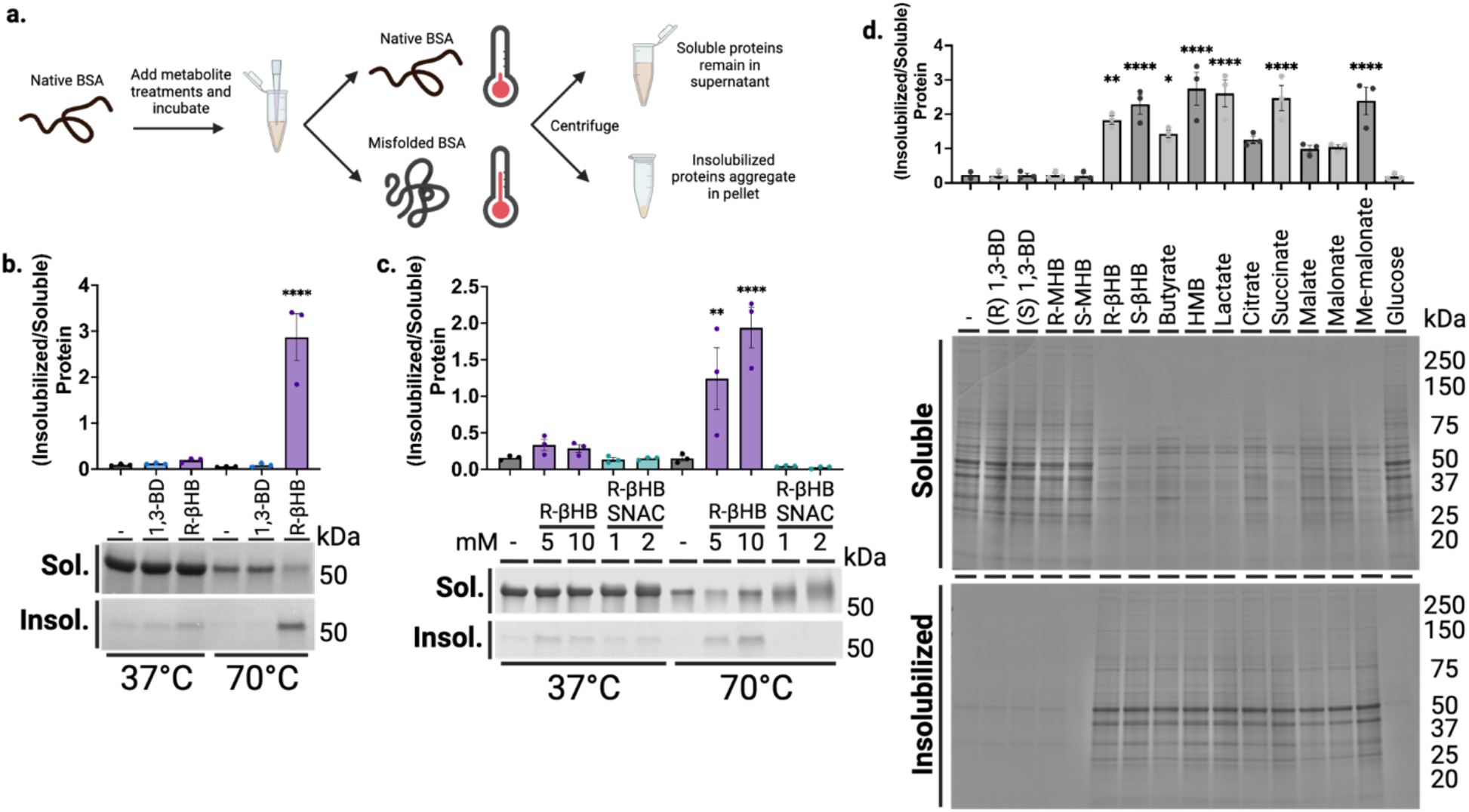
β-hydroxybutyrate directly induces protein insolubility without posttranslational modification. **a,** Schematic of experimental procedures. **b-c,** Imperial staining and quantification of soluble and induced-insoluble native (+37°C) and heat-misfolded (+70°C) BSA treated with **(b)** 10 mM of R-βHB or 10 mM of alcohol analogue, 1,3-BD, and **(c)** 5-10 mM of R-βHB or 1-2 mM of nonenzymatic elicitor of KβHB, (R)-βHB-SNAC. **d,** Imperial staining and quantification of 24 month male wild-type mouse soluble cytosolic brain proteins which remain soluble or are insolubilized after treatment with 10 mM of a small library of 15 metabolites. b, Mean ± S.E.M, N=3, p-value calculated using one-way ANOVA with Sidak’s multiple comparison test. c, Mean ± S.E.M, N=3, p-value calculated using one-way ANOVA with Sidak’s multiple comparison test. d, Mean ± S.E.M, N=3, p-value calculated using one-way ANOVA with Dunnett’s multiple comparison test.

R-βHB is substrate for the posttranslational modification lysine β-hydroxybutyrylation (KβHB) on both histone and non-histone proteins^22^, which could potentially affect protein folding and solubility. We used an *in vitro* mimetic of R-βHB-CoA, R-βHB-SNAC, to non-enzymatically induce KβHB and directly test if KβHB mediates misfolded protein insolubility. R-βHB-SNAC and not R-βHB, as measured by immunoblotting for KβHB, induces KβHB on heat-misfolded BSA (Extended Data 2b). However, we found that R-βHB-SNAC cannot induce insolubility of heat-misfolded BSA, as measured by Imperial staining (Fig. 1c).

To identify the response of misfolded proteins to simultaneous insolubilization and posttranslational modification pressures, we co-administered R-βHB (5-10 mM) and R-βHB-SNAC (0.5-2 mM) during heat-induced misfolding of BSA, measured by Imperial staining (Extended Data 2c). We observed that 2 mM R-βHB-SNAC attenuated the induction of heat-misfolded BSA insolubility by 10 mM R-βHB, and R-βHB-induced insolubility at 5 mM was effectively abrogated by all R-βHB-SNAC concentrations. To further test whether R-βHB and KβHB interact on similar sites on BSA, and which interaction was dominant, we tested whether pretreatment of BSA with R-βHB-SNAC would alter R-βHB-induced insolubility. We completed two experiments, with either native folded KβHB-BSA or heat-misfolded KβHB-BSA. Firstly, we induced native folded KβHB-BSA by incubating R-βHB-SNAC (0.5-2 mM) with BSA at +37°C, then exposed KβHB-BSA to heat-misfolding and R-βHB treatment (5-10 mM) after filtering to remove R-βHB-SNAC (Extended Data 2d). Secondly, as heat misfolding may expose previously inaccessible lysine residues, we induced KβHB on heat-misfolded BSA by incubating increasing concentrations of R-βHB-SNAC with BSA at +70°C, then exposed this KβHB-BSA to R-βHB treatment (5-10 mM) after filtering to remove R-βHB-SNAC (Extended Data 2e). In both instances, 10 mM R-βHB induced insolubility of heat-misfolded KβHB- BSA, regardless of whether KβHB was elicited on native folded or heat-misfolded KβHB-BSA, or the concentration of R-βHB-SNAC used. Together, these data show that R-βHB-induced insolubility does not occur via KβHB, that KβHB and R-βHB have opposing effects on the solubility of misfolded proteins, and that the effect of R-βHB is dominant over KβHB.

After establishing and validating βHB-induced insolubilization of heat-misfolded purified proteins, we tested the insolubilization effect in a heterogenous mix of relevant protein targets, mouse brain lysate. We chose to examine an aged brain environment to test R-βHB-induced insolubility in the relevant setting of soluble misfolded proteins that accumulate throughout aging. Additionally, we sought to examine if other metabolites structurally similar to R-βHB could induce insolubility (Extended Data 1c). We extracted and homogenized whole brains from 24 month wild-type (C57BL/6) male mice and used subcellular fractionation to isolate soluble cytosolic proteins from the homogenate (Extended Data 3a). We performed an *ex vivo* insolubilization assay at +37°C, by incubating buffered soluble cytosolic proteins with 10 mM of R- and S-1,3-BD, R- and S-methyl-hydroxybutyrate (MHB), and R- and S-βHB, as well as butyrate, hydroxymethylbutyrate (HMB), lactate, citrate, succinate, malate, malonate, methyl-malonate, and glucose, and used centrifugation to pellet proteins which became insolubilized after incubation, quantified by Imperial staining (Fig. 1d). We found that some, but not all metabolites, including R- and S-βHB, butyrate, HMB, lactate, succinate, and methyl-malonate significantly insolubilized previously-soluble proteins. Metabolites that induced protein insolubilization shared a common carboxylic acid structural feature and insolubilized a wide variety of protein bands, with affinity for high-molecular weight proteins. While other metabolites may possess similar or stronger protein insolubilization properties than R-βHB, we continued to focus on R-βHB due to the high dynamic range of physiological concentrations, relevance to ongoing clinical investigations of metabolic therapies for NDDs, and the existance of a diverse set of ketone body-elevating experimental tools, some of which are used below.

### *Ex vivo* identification of (R)-β-hydroxybutyrate insolubilization targets across the mouse brain proteome

We next used the *ex vivo* insolubilization assay, coupled with mass spectrometry proteomics, to identify a library of dose-dependent R-βHB targets from male mouse brains (Fig. 2a). First, we tested R-βHB (1-10 mM), compared to 1,3-BD (1-10 mM), in 24 month wild-type brain lysate measured by Imperial staining (Fig. 2b). The significant insolubilization at 5 and 10 mM R-βHB involved the deposition of proteins from a wide range of molecular weights. We confirmed these results were similar in age- and strain-matched female mouse brain lysate, and proceeded with male tissue for subsequent experiments (Extended Data 3b). We selected the 1 and 5 mM R-βHB conditions for proteomics analysis, as these concentrations represent the lower and upper plasma concentration thresholds of physiological ketosis achieved with fasting, standard ketogenic diets, and most exogenous ketones in mammals^14^. 10 mM R-βHB is commonly utilized experimentally *in vitro* to elicit maximal relevant cellular effects^24, 25^. For our coupled *ex vivo* insolubilization assay and proteomics experiment, we hypothesized that 1 mM R-βHB would identify high-affinity direct targets, while 5 mM R-βHB would reveal lower-affinity and indirect targets. We analyzed the protein pellets treated with 0, 1, and 5 mM R-βHB by data-independent acquisition mass spectrometry (DIA-MS), with 0 mM R-βHB serving as the reference^58, 59^. Our downstream analysis assessed the differential protein enrichment of the 1/0 mM R-βHB and 5/0 mM R-βHB groups. We found that each treatment group clustered separately when examined with partial least squares-discriminant analysis (PLS-DA) (Fig. 2c). 3,283 proteins comprised the detectable proteome across both comparison groups. The 791 significantly regulated proteins in the 1/0 mM R-βHB comparison were largely enriched in the 1 mM direction, visualized with a volcano plot (Fig. 2d). The 3,232 significantly regulated proteins in the 5/0 mM R-βHB were largely enriched in the 5 mM direction, visualized with a volcano plot (Fig. 2e). 790 of the 791 proteins from the 1/0 mM R-βHB group were represented in the 5/0 mM R-βHB group, visualized with a Venn diagram (Fig. 2f). Overall, we found that R-βHB interacted with a large fraction of the mouse brain proteome and almost exclusively increased protein insolubility.

**Fig. 2.**
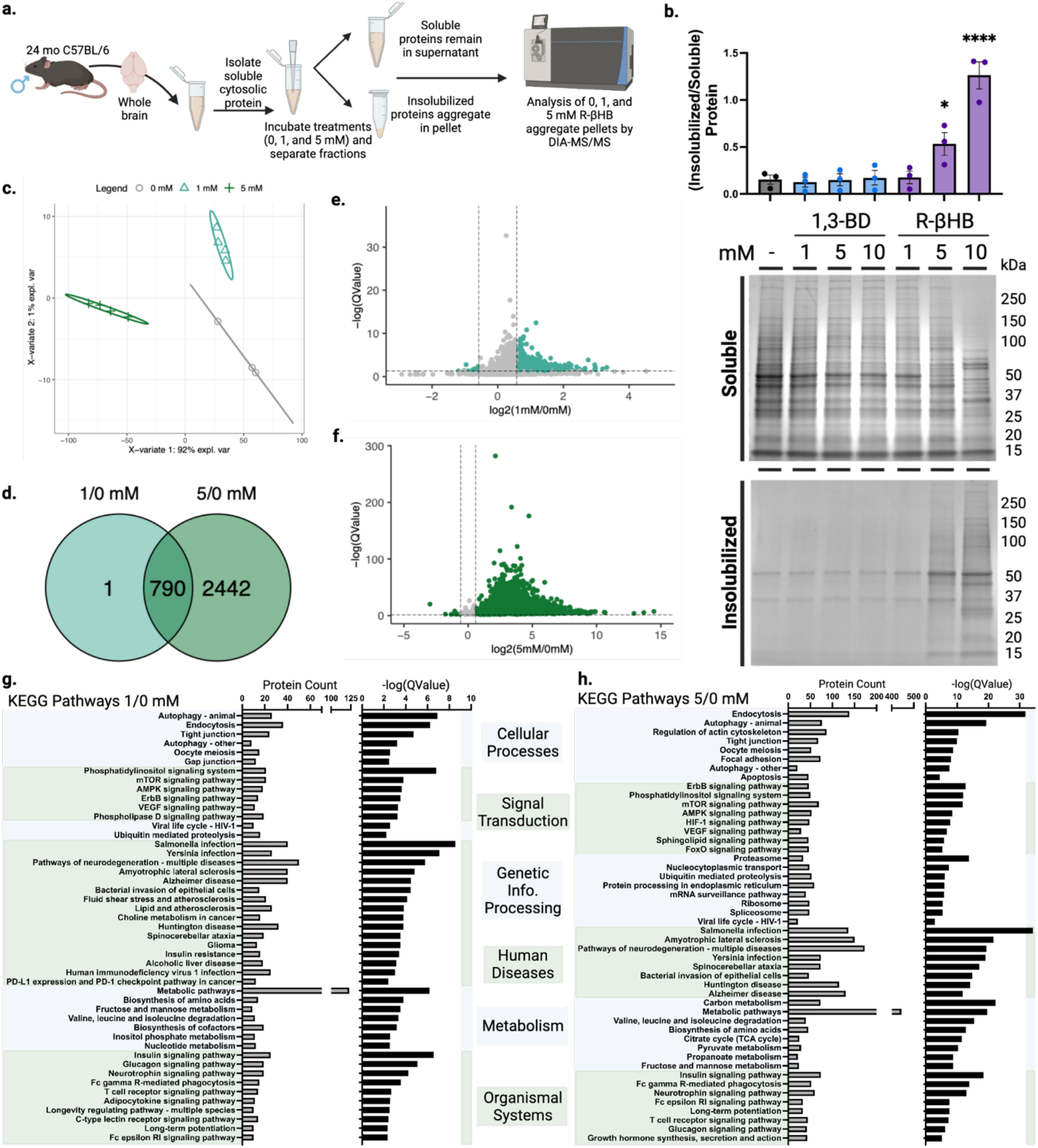
β-hydroxybutyrate remodels the older C57BL/6 mouse brain proteome *ex vivo* via insolubilization of targets. **a,** Schematic of experimental procedures. **b,** Imperial staining and quantification of 24 month male wild-type mouse soluble cytosolic brain proteins which remain soluble or are insolubilized after treatment with 1-10 mM of R-βHB and 1-10 mM of 1,3-BD. **c,** Partial least squares-discriminant analysis (PLS-DA) of clustering of 0, 1, and 5 mM R-βHB proteomic samples. **d,** Venn diagram comparing target proteins in 1/0 mM and 5/0 mM R-βHB proteomics samples. **e-f,** Volcano plots of **(e)** 1/0 mM and **(f)** 5/0 mM R-βHB proteomic samples. **g-h,** Results from clusterProfiler KEGG overrepresentation analysis with BRITE functional hierarchical classifications, arranged by decreasing −log(QValue) for **(g)** 1/0 mM group, all KEGG pathways shown, and **(h)** 5/0 mM group, top 8 KEGG pathways of each BRITE category shown. b, Mean ± S.E.M, N=3, p-value calculated using one-way ANOVA with Dunnett’s multiple comparison test.

Gene ontology overrepresentation analysis for biological process on the proteins significantly insolubilized by R-βHB revealed that proteins in both 1/0 mM and 5/0 mM treatment groups were significantly associated with cellular protein metabolic processes, cellular localization, protein localization, organelle organization, establishment of localization in cell, and cellular catabolism (Extended Data 3d-e). KEGG pathway overrepresentation analysis on the proteins significantly deposited by R-βHB identified significantly regulated KEGG pathways which fell within all possible BRITE hierarchical categories, with all pathways shown for 1/0 mM R-βHB (Fig. 2g) and top 8 from each category shown for 5/0 mM R-βHB (Fig. 2h). Both overrepresentation analyses used the whole mouse genome as a background, sourced from the R package org.Mm.eg.db, were completed using ClusterProfiler in R, and are ranked by associated −log(QValue). Within the BRITE category “Human Disease”, multiple pathways associated with neurodegenerative disease were identified, including AD and Huntington Disease (HD), Amyotrophic Lateral Sclerosis (ALS), and Multiple Pathways of Neurodegeneration. Additionally, protein degradation pathways such as Autophagy, Proteasome, and Ubiquitin Mediated Proteolysis were also identified. The only subcategorization within the BRITE category “Environmental Information Processing” was “Signal Transduction” for both treatment groups, including aging-related mTOR, AMPK, and FOXO signaling pathways. While wild-type mice do not experience the pathology of NDDs exactly like humans, the selectivity of R-βHB-induced insolubilization for proteins within NDD pathways and protein degradation mechanisms suggests that this mechanism may be neuroprotective, which we sought to test directly. We hypothesized that R-βHB-induced insolubilization assists in the clearance of misfolded, toxic, and disease-associated proteins in order to promote efficiency of a system under metabolic stress.

### β-hydroxybutyrate induces structural remodeling of proteins

Having previously observed that R-βHB alters the conformation of heat-misfolded albumin to decrease nonpolar cavities, we next hypothesized that the R-βHB-induced insolubilization of proteins decreased the cellular toxicity of these proteins. To test whether βHB-induced insolubilization could directly interact with pathogenic protein structure and conformation, we monitored β-sheet content of native brain soluble cytosolic protein derived from 24 month wild-type mouse incubated with R-βHB (1-10 mM) (Fig. 3a-b, Extended Data 4a-b). We tested R-βHB in these lysates using a modified protocol from our *ex vivo* insolubilization assay, incubating within a plate reader to assess thioflavin T fluorescence. Increased thioflavin T fluorescence corresponds to increased β-sheet content within proteins and is used to monitor protein oligomerization and fibrillization, especially of amyloid-β and other NDD-related proteins. In males, we identified significant decreases in β-sheet content, measured by area under the curve, within soluble cytosolic proteins treated with 2-10 mM R-βHB, despite the increased insolubilization shown in our *ex vivo* insolubilization assay (Fig. 3b). We observed a similar decrease in β-sheet content using female brain lysates (Extended Data 4b). These data show that R-βHB directly alters the conformation of proteins targeted for insolubilization.

**Fig. 3.**
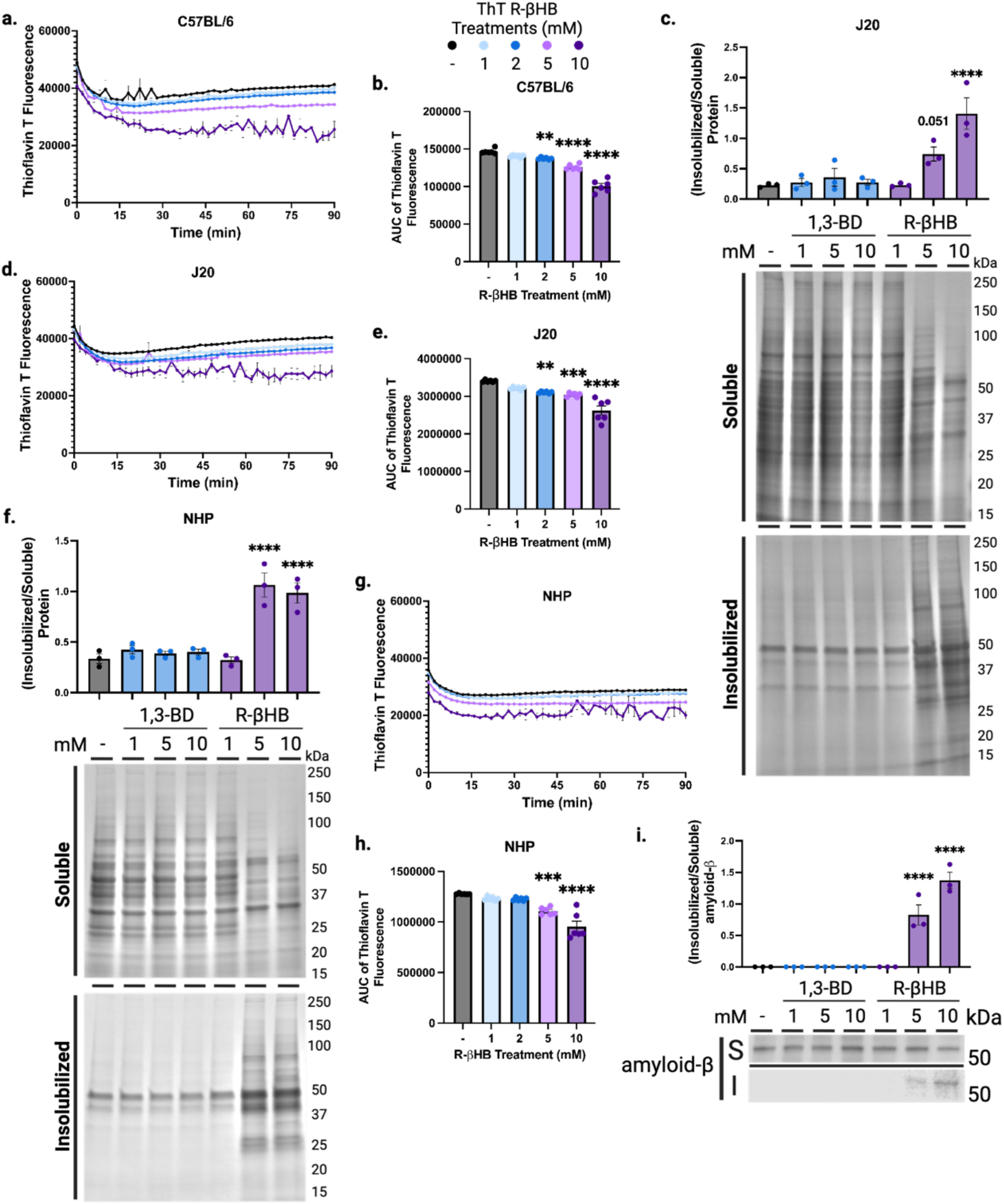
β-hydroxybutyrate induces structural remodeling of proteins. **a-b,** Total protein aggregation kinetics monitored via Thioflavin T fluorescence in mouse soluble cytosolic brain proteins treated with 1-10 mM of R-βHB from 24 month wild-type male, **(a)** timecourse and **(b)** area under the curve. **c,** Imperial stain of 4 month male J20 mouse soluble cytosolic brain proteins which remain soluble or are insolubilized after treatment with 1-10 mM of R-βHB and 1-10 mM of 1,3-BD. **d-e,** Total protein aggregation kinetics monitored via Thioflavin T fluorescence in mouse soluble cytosolic brain proteins treated with 1-10 mM of R-βHB from 4 month J20 male, **(d)** timecourse and **(e)** area under the curve. **f,** Imperial stain of 26 year male non-human primate soluble cytosolic brain proteins which remain soluble or are insolubilized after treatment with 1-10 mM of R-βHB and 1-10 mM of 1,3-BD**. g-h,** Total protein aggregation kinetics monitored via Thioflavin T fluorescence in non-human primate soluble cytosolic brain proteins treated with 1-10 mM of R-βHB from 26 year rhesus macaque male, **(g)** timecourse and **(h)** area under the curve. **i,** Western blot and quantification for amyloid-β (6E10) in soluble cytosolic J20 mouse brain proteins which remain soluble or are insolubilized after treatment with 1-10 mM of R-βHB and 1-10 mM of 1,3-BD. a, Mean ± S.E.M, N=6, p-value calculated using two-way ANOVA with Dunnett’s multiple comparison test. b, Mean ± S.E.M, N=6, p-value calculated using one-way ANOVA with Dunnett’s multiple comparison test. c, Mean ± S.E.M, N=3, p-value calculated using one-way ANOVA with Dunnett’s multiple comparison test. d, Mean ± S.E.M, N=6, p-value calculated using one-way ANOVA with Dunnett’s multiple comparison test. e, Mean ± S.E.M, N=6, p-value calculated using one-way ANOVA with Dunnett’s multiple comparison test. f, Mean ± S.E.M, N=3, p-value calculated using one-way ANOVA with Dunnett’s multiple comparison test. g, Mean ± S.E.M, N=6, p-value calculated using one-way ANOVA with Dunnett’s multiple comparison test. h, Mean ± S.E.M, N=6, p-value calculated using one-way ANOVA with Dunnett’s multiple comparison test. i, Mean ± S.E.M, N=3, p-value calculated using one-way ANOVA with Dunnett’s multiple comparison test.

Based on our overrepresentation analyses and the direct interactions with protein conformation, we chose to further investigate the interactions between R-βHB and amyloid-β by using brain tissue from 4 month J20 mice (which express human APP with AD-related genetic mutations). An *ex vivo* insolubilization assay with 4 month male J20 mouse lysate by R-βHB and 1,3-BD matched the insolubilization pattern from 24 month wild-type mice, quantified by Imperial staining (Fig. 3c). We additionally monitored β-sheet content of native brain soluble cytosolic protein derived from 4 month J20 mouse incubated with R-βHB (1-10 mM) (Fig. 3d-e). Here, we uncovered a similar trend of significant decreases in β-sheet content within soluble cytosolic proteins treated with 2-10 mM R-βHB (Fig. 3e).

To further examine the effects of R-βHB in a brain environment more similar to humans, we utilized brain tissue from male and female aged *Macaca mulatta* (Rhesus Macaque). The *ex vivo* insolubilization assay with 26 year male brain lysate by R-βHB and 1,3-BD matched the insolubilization data from older wild-type mice, with higher induction of insolubilization at 5 mM R-βHB in *M. mulatta* than in mouse (Fig. 3f). We replicated our assay of β-sheet content with native brain soluble cytosolic protein derived from 26 year male *M. mulatta* incubated with R-βHB (1-10 mM) (Fig. 3g-h). Area under the curve quantification matched mouse, with significant decreases in β-sheet content within soluble cytosolic proteins treated with 5-10 mM R-βHB (Fig. 3h).

In 25 year *M. mulatta* female brain lysate, insolubilization patterns matched mouse and male *M. mulatta* data (Extended Data 4c). We assayed β-sheet content kinetics with native brain soluble cytosolic protein derived from 25 year female *M. mulatta* incubated with R-βHB (1-10 mM) (Extended Data 4d-e). Area under the curve quantification matched mouse and male *M. mulatta* results as well, with significant decreases in β-sheet content within soluble cytosolic proteins treated with 5-10 mM R-βHB (Extended Data 4e). Overall, these data show the capability of R-βHB to induce structural changes in proteins in both the mouse and non-human primate brain to ultimately lower structural conformations associated with NDDs, suggesting a positive role for R-βHB in proteostasis.

### β-hydroxybutyrate inhibits amyloid-β aggregation and oligomeric toxicity

Thus far, our data revealed that R-βHB-induced insolubilization targets in older wild-type mice included proteins related to AD, that *ex vivo* insolubilization data in J20 mouse brain tissue matched older wild-type mouse, and that R-βHB-induced insolubilization equates to decreased β-sheet content in both mouse and non-human primate brain tissue. We next sought to clarify specific effects of R-βHB-induced insolubilization on the AD-related pathogenic protein amyloid-β. Firstly, we identified an insolubilization effect on oligomeric amyloid-β in J20 mouse brain tissue, visualized through immunoblotting for amyloid-β_1-16_ post-*ex vivo* insolubilization assay (Fig. 3i). Here, we identified a soluble amyloid-β oligomer structure of roughly 50 kD which was significantly insolubilized by 10 mM R-βHB. Furthermore, incubation of a fluorescent amyloid-β_1-42_ peptide with native soluble cytosolic proteins from 24 month wild-type mice led to development of a soluble oligomeric smear, which was insolubilized by R-βHB (Fig. 4a).

**Fig. 4.**
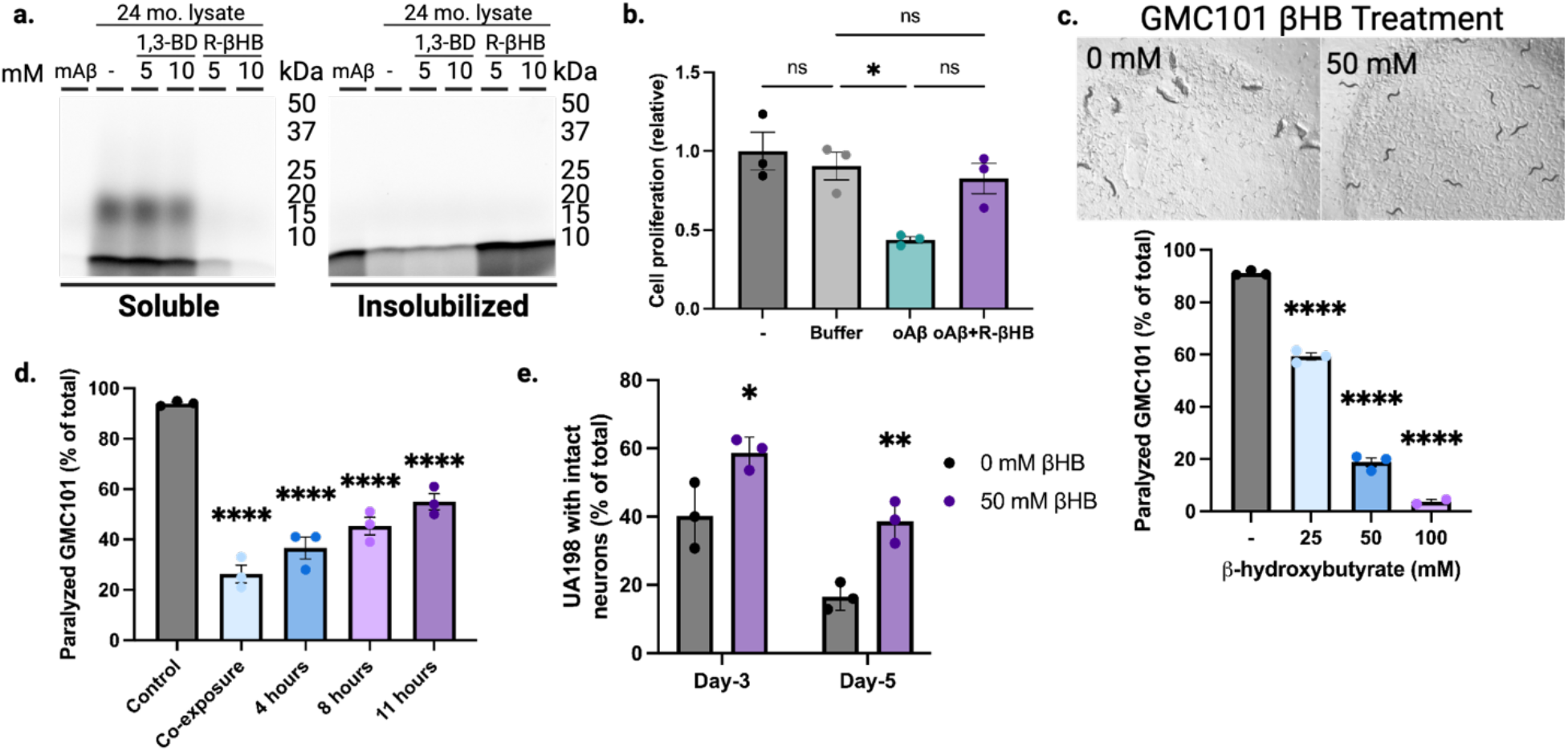
| β-hydroxybutyrate inhibits oligomeric toxicity and suppresses human amyloid-β-induced paralysis and neurotoxicity in *C. elegans*. **a,** SDS-PAGE of HiLyte Fluor 488-labeled amyloid-β_1-42_ peptide monomer (mAβ) incubated with 24 month male wild-type soluble cytosolic brain proteins and treated with 5-10 mM of R-βHB and 5-10 mM of 1,3-BD. **b,** Quantification of N2a cell proliferation monitored by XTT Assay following treatment with 2 µM amyloid-β oligomers (oAβ) and 10 mM R-βHB. **c-d,** Quantification of amyloid-β proteotoxicity in temperature-sensitive (aggregation-permissive at +25°C) GMC101 strain, determined by scoring the percentage of animals paralyzed at **(c)** 25-28 hours following temperature shift and with 25-100 mM of βHB treatment (representative image shown), and **(d)** following temperature shift without treatment, then movement to 50 mM βHB treatment at varying timepoints. **e,** Quantification of amyloid-β neurotoxicity was determined by scoring number of intact glutaminergic neurons in UA198 animals (expressing amyloid-β in GFP-tagged glutaminergic neurons) with 50 mM of βHB treatment. a, Representative image from triplicate repetitions. b, Mean ± S.E.M, N=3, p-value calculated using one-way ANOVA with Tukey’s multiple comparison test. c-d, Mean ± S.E.M, N=3 (∼300 animals), p-value calculated using one-way ANOVA with Dunnett’s multiple comparison test. e, Mean ± S.E.M, N=3 (∼300 animals). p-value calculated using two-way ANOVA with Sidak’s multiple comparison test.

To test whether these structural and solubility changes altered cytotoxicity, we incubated N2a mouse neuroblastoma neuronal cells with 2 µM amyloid-β oligomers with or without 10 mM R-βHB (Fig. 4b). Compared to control, cell viability measured by XTT assay significantly decreased after 24 hours of 2 µM amyloid-β oligomers, but was ameliorated by the addition of 10 mM R-βHB. Therefore, R-βHB-induced insolubilization interferes with the oligomerization of amyloid-β peptides and insolubilizes both oligomeric and high-molecular weight structures of pathogenic proteins, ultimately inhibiting oligomeric cytotoxicity in cell culture.

### β-hydroxybutyrate suppresses human amyloid-β-induced paralysis and neurotoxicity in *C. elegans*

To test whether βHB-induced insolubility reduces amyloid-β toxicity at an organismal level, we chose amyloid-β_1-42_ overexpressing models of the nematode *Caenorhabditis elegans*. We chose these models for the specificity of expressing amyloid-β_1-42_ in specific tissues with direct functional phenotypes. If βHB-induced insolubility enhanced amyloid-β pathogenicity via increased aggregation, it would be detrimental in these models. Firstly, we confirmed that there was no effect of βHB or R-βHB on bacterial growth, when compounds were added to the bacterial feeding solution covering culture plates (Extended Data 4f). Next, we quantified proteotoxicity-induced paralysis in GMC101 animals (expressing temperature-sensitive human amyloid-β_1-42_ in body wall muscle cells) following shift to aggregation-permissive +25°C and transfer to plates covered with bacterial feeding solution containing βHB (25-100 mM) (Fig. 4c with representative image at 50 mM βHB). Animals were scored 25-28 hours post-temperature shift. We identified a significant decrease in the percentage of animals experiencing proteotoxicity-induced paralysis at all βHB concentrations, with 100 mM nearly completely rescuing paralysis. We next tested the ability of βHB to suppress the proteotoxic paralysis phenotype following a period of aggregation-permissive incubation. GMC101 animals were all shifted to +25°C on plates lacking βHB, then after 0-11 hours were transferred to new plates with 50 mM βHB (Fig. 4d). βHB rescued paralysis at all timepoints, compared to control, even when treatment began 11 hours after shift to the aggregation-permissive temperature. The robust suppresion of amyloid-β_1-42_ aggregation-induced paralysis in this model supports a positive functional role for βHB-induced insolubility.

Next, we used the UA198 strain (expressing human amyloid-β_1-42_ in GFP-labeled glutaminergic neurons) to model amyloid-β neurotoxicity. C. elegans have 5 glutaminergic neurons in the tail region which experience age-related degeneration exacerbated by amyloid-β_1-42_ aggregation. We transferred UA198 animals to plates covered with 50 mM βHB and scored the number of intact neurons with fluorescence microscopy at day-3 and day-5 (Fig. 4e). At both timepoints, βHB-treated animals retained significantly higher proportions of intact neurons, with day-5 βHB-treated animals having equivalent intact neurons as day-3 control animals. The preservation of intact neurons in older βHB-treated animals corroborates our increased mouse neuronal cell viability *in vitro* and further supports a positive functional impact of βHB-induced insolubility in the brain.

### Subchronic treatment with a ketone ester remodels the older C57BL/6 brain insolublome *in vivo*

Thus far, we had characterized R-βHB-induced insolubility when added *ex vivo* to protein extracts. To test the physiological relevance of this biochemical activity, and address any potential confounding *in vitro* mechanisms, we tested whether R-βHB- induced insolubility could be observed in a fully *in vivo* system. To date, multiple fasting-metabolism interventions which elevate ketone body concentrations in the blood and brain have been shown to promote clearance of pathological proteins in the brain^31, 41–45^. However, no studies have identified the intermediate insolubilization mechanism we have uncovered *in vitro* and *ex vivo*. Ketone esters are the most efficient pharmacological delivery method for elevating ketone bodies in both blood and brain, and can do so without dietary changes. Bis hexanoyl (R)-1,3-butanediol (BH-BD) is a ketone ester comprised of 1,3-butanediol and medium-chain fatty acid moieties which is cleaved in the small intestine, with its constituents undergoing rapid metabolism in the liver to produce ketone bodies for export to extrahepatic organs^60, 61^ (Extended Data 1c). We verified with mass spectrometry that BH-BD increased R-βHB concentrations in the 22 month wild-type mouse brain following a 7 day feeding schedule, compared to a control diet (CD) (Extended Data 5a). We next hypothesized that subchronic treatment with BH-BD in older mice would remodel the brain native insoluble protein compartment (insolublome).

To test this hypothesis, we administered 5g/kg BH-BD or isocaloric canola oil, as a non-ketogenic control, via oral gavage to 20 month wild-type male mice twice daily for 7 days and collected blood by tail-bleed each evening, 1 hour post-gavage (Extended Data 5b). We observed no significant changes in body weight between cohorts (Extended Data 5c). BH-BD elicited a significant decrease in plasma glucose compared to control (Extended Data 5d). Additionally, BH-BD induced ketosis in the mice, significantly elevating plasma βHB and peaking above 4 mM on the third day (Extended Data 5e).

After the end of 7 days, we harvested tissue and used subcellular fractionation to separate the native TEM soluble and insoluble compartments of the aged brain. To more rigorously examine the subtle changes in the brain insolublome with finer detail, we employed a sequential detergent extraction to fractionate the insolublome by detergent resistance into four compartments (Fig. 5a). Each fraction contains proteins which are more resistant to increasingly aggressive detergent buffers. The initial native insoluble protein pellet was resuspended in TEM + 0.5% NP40. The resolubilized solution was incubated at +37°C and centrifuged to produce fraction 1 (F1) in the supernatant. The resulting pellet was resuspended in TEM + 0.5% NP40 + 0.5% Sodium Deoxycholate + 0.25% SDS, incubated at +37°C, and centrifuged to produce the supernatant fraction 2 (F2) and a pellet. This pellet was resuspended in TEM + 0.5% NP40 + 0.5% Sodium Deoxycholate + 2% SDS. The resolubilized solution was incubated at +37°C and centrifuged to produce the supernatant fraction 3 (F3) and a final pellet. This final pellet was resuspended in TEM + 0.5% NP40 + 0.5% Sodium Deoxycholate + 3% SDS. The resolubilized solution was incubated at +25°C overnight before centrifugation to produce the final supernatant fraction 4 (F4).

**Fig. 5.**
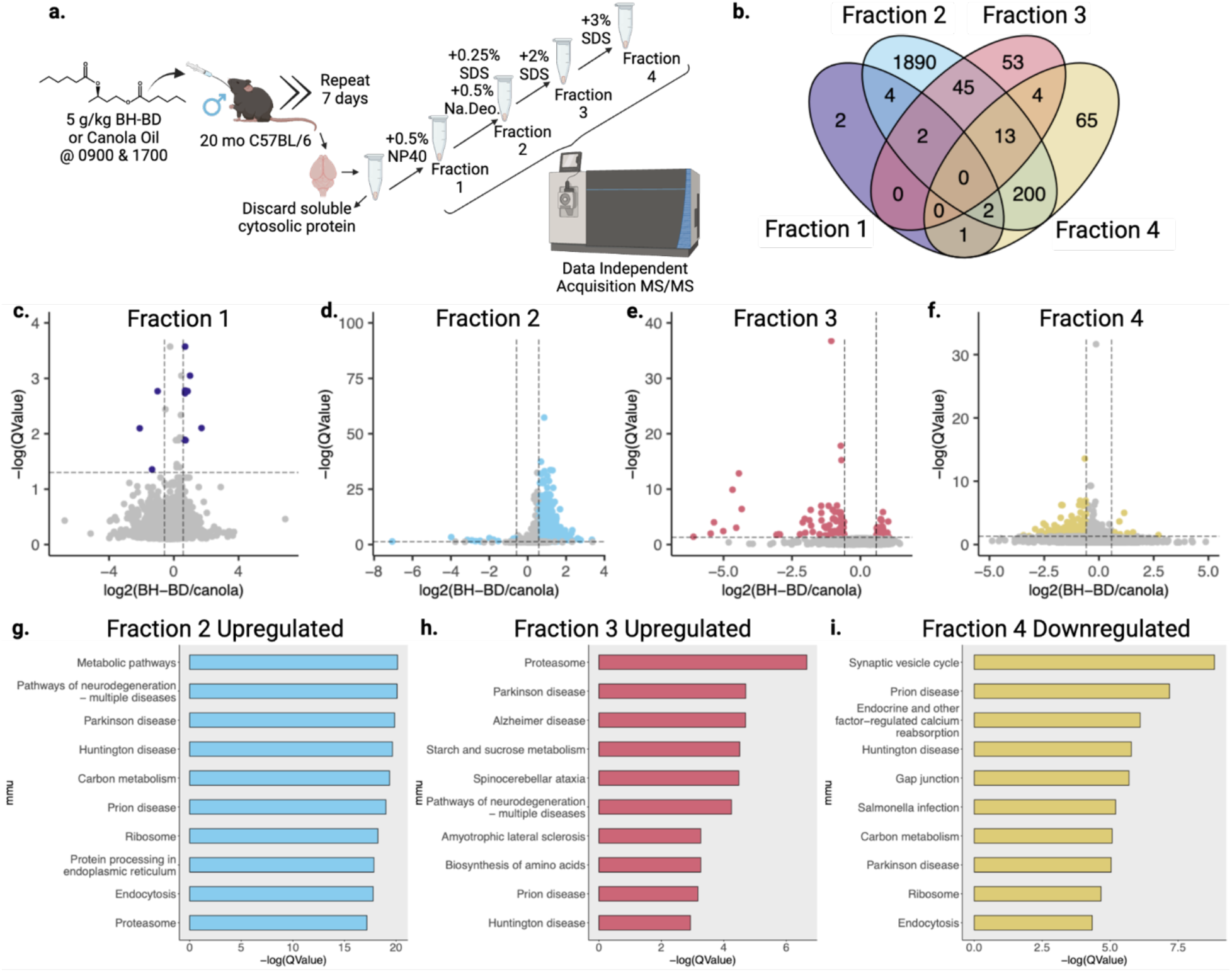
Subchronic treatment with a ketone ester remodels the older C57BL/6 brain insolublome *in vivo*. **a,** Schematic of cohort and sequential detergent extraction used on insoluble cytosolic brain proteins for proteomic analysis. **b,** Venn diagram of significantly regulated proteins with ketone ester BH-BD/canola oil control supplementation from each detergent fraction. **c-f,** Volcano plots of BH-BD/control proteomic samples from each detergent fraction. **g-i,** Dotplots from clusterProfiler KEGG overrepresentation analysis on **(g)** upregulated proteins in fraction 2, **(h)** upregulated proteins in fraction 3, and **(i)** downregulated proteins in fraction 4.

Equal mass of protein from each treatment group of the four fractions was analyzed by DIA-MS. All four fractions clustered separately when examined with PLS-DA (Extended Data 6b-e). The detectable proteome across all four fractions varied from a minimum of 2,693 to a maximum of 3,118 proteins detected. Our downstream analysis assessed the differential protein abundance of the BH-BD/canola insolublome in each of the four fractions, with “upregulated” indicating proteins with higher abundance (i.e. higher insolubility) in the BH-BD group and “downregulated” indicating higher abundance in the control canola group. The significantly regulated proteins in each of the four fractions are distinct, though some overlap between fractions occur, as visualized by Venn diagram (Fig. 5b). F2 is the most distinct of the four fractions, with the majority of proteins unshared with any other fraction. The 11 significantly regulated proteins in F1 favored upregulation, visualized with a volcano plot (Fig. 5c). In F2, the 2156 significantly regulated proteins also favored upregulation, visualized with a volcano plot (Fig. 5d). In F3, the 117 significantly regulated proteins instead favored downregulation, visualized with a volcano plot (Fig. 5e). Finally, in F4, the 285 significantly regulated proteins also favored downregulation, visualized with a volcano plot (Fig. 5f). This pattern suggested that R-βHB targets were being increasingly insolubilized in lower and medium insoluble states, but cleared from highly insoluble states.

We analyzed patterns in the BH-BD-remodeled brain insolublome using Gene ontology and KEGG. Gene ontology overrepresentation analysis showed distinct pathways enriched in each fraction (Extended Data 7). KEGG overrepresentation analysis on upregulated and downregulated proteins in each fraction showed a dramatic remodeling of the brain insolublome following BH-BD treatment, compared to control (Fig. 5g-i). We observed no enrichment for downregulated proteins in F1, and found that upregulated proteins in F1 favored pathways related to metabolism (Extended Data 7a). In F2, we found that upregulated proteins were highly associated with NDDs and the proteasome, unlike downregulated proteins in F2 (Fig. 4g, Extended Data 7b). In F3, we similarly found a strong association with upregulated proteins and NDDs, including AD and the proteasome, with downregulated proteins showing no associations with NDDs (Fig. 4h, Extended Data 7c). Finally, in F4, we observed that downregulated proteins were highly associated with synaptic vesicle cycle and NDDs (Fig. 5i). Together, these data suggest that the control insolublome models aging, with proteins related to NDDs settling into the most insoluble compartment of the insolublome. Following BH-BD administration, the insolublome is completely remodeled, with medium insoluble fractions F2 and F3 demonstrating βHB-induced insolubilization of NDD-related proteins, and highly insoluble fraction F4 displaying clearance of these NDD-related proteins.

### β-hydroxybutyrate targets display common structural features and are cleared through protein degradation pathways

To further dissect the relationships between βHB-targeted proteins and to identify core structural sequences that transcend protein functional pathways, we examined the prevalence of InterPro protein domains within each DIA-MS group. In the *ex vivo* protein insolubilization assay groups, we pinpointed a distinct shift in significantly regulated protein domains from 1/0 mM R-βHB to 5/0 mM R-βHB treatment groups. We found that tubulin-related domains comprised 8 of the top 10 most significantly enriched protein domains of the proteins significantly deposited by R-βHB in the 1/0 mM R-βHB treatment group (Fig. 6a). Conversely, in the 5/0 mM R-βHB treatment group, we found that significantly enrichment protein domains included many related to cellular signaling, including protein-protein interactions, such as ARM-like, ARM-type fold, WD40 repeat, WD40 repeat domain superfamily, and PH-like domain superfamily, as well as domains related to NAD, GTP, and NTPase activity (Fig. 6b). The only common top 10 significantly enriched protein domain shared between 1/0 mM R-βHB to 5/0 mM R-βHB treatment groups was the Rossmann-like α/β/α fold.

**Fig. 6.**
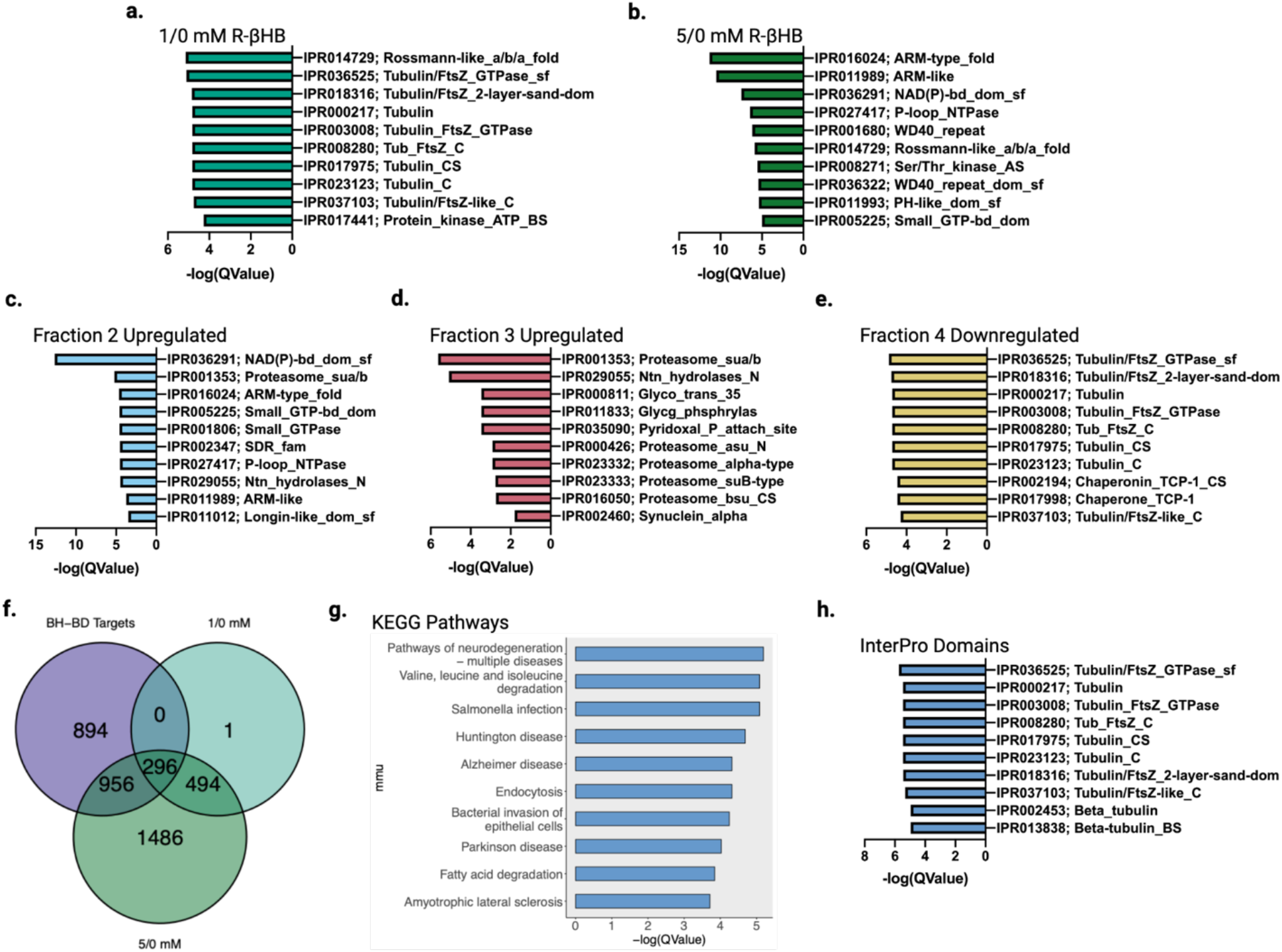
β-hydroxybutyrate targets display common structural features and are cleared through protein degradation pathways. **a-e,** Top 10 significantly enriched protein domains ranked by Q-value from **(a)** 1/0 mM R-βHB *ex vivo* upregulated proteins, **(b)** 5/0 mM R-βHB *ex vivo* upregulated proteins, **(c)** BH-BD/control fraction 2 upregulated proteins, **(d)** BH-BD/control fraction 3 upregulated proteins, and **(e)** BH-BD/control fraction 4 downregulated proteins. **f,** Venn diagram of BH-BD/control significantly upregulated proteins from all fractions crossed with upregulated proteins from 1/0 mM R-βHB and 5/0 mM R-βHB *ex vivo* proteomics groups, the 296 primary protein targets of R-βHB were calculated to have a p-value of 0.000001. **g,** Dotplot from clusterProfiler KEGG overrepresentation analysis on 296 primary protein targets of R-βHB from Fig. 6f. **h,** Top 10 significantly enriched protein domains ranked by Q-value from 296 primary protein targets of R-βHB from Fig. 6f. f, p-value calculated using one-tailed probability test giving a z score = −54.4.

Protein domains identified in the *in vivo* BH-BD/canola insolublome fractions connected larger biological themes examined in the *ex vivo* R-βHB protein domain targets and the protein functional pathways from both *ex vivo* and *in vivo* KEGG overrepresentation analysis. Interestingly, it appears that *in vivo* F2 and F3 proteins seem to mirror the *ex vivo* 5/0 mM R-βHB treatment group, while *in vivo* F4 proteins seem to mirror the ex vivo 1/0 mM R-βHB treatment group. *In vivo* F2 upregulated proteins share 5 of their top 10 significantly enriched protein domains with the *ex vivo* 5/0 mM R-βHB treatment group, on top of including domains related to NAD activity and the proteasome (Fig. 6c). F3 upregulated proteins, mirroring functional pathway themes identified with KEGG overrepresentation analysis, display proteasome-related domains in 5 of the top 10 significantly enriched (Fig. 6d). Additionally, the domain α-synuclein, a protein heavily dysregulated in the NDD, Parkinson disease (PD), was enriched in F3 upregulated proteins. Indeed, the relationship of *ex vivo* and *in vivo* becomes clear in the F4 downregulated proteins, targets which have been retained in the canola-group brains but cleared from BH-BD-group brains. Here, the same 8 tubulin-related domains from 1/0 mM R-βHB are again in the top 10 significantly enriched domains for downregulated F4 proteins (Fig. 6e). Top 10 significantly enriched protein domains from F3 downregulated proteins showed similarities with gene ontology for biological process (Extended Data 8a).

To identify the primary targets of R-βHB, we overlapped the proteins upregulated in all four *in vivo* fractions and proteins insolubilized with *ex vivo* 1/0 mM and 5/0 mM R-βHB treatment (Fig. 6f). 296 proteins were shared between these groups. We analyzed functional pathways in these primary R-βHB targets using KEGG analysis. KEGG overrepresentation analysis identified Multiple Pathways of Neurodegeneration as the most significantly regulated identifier of the primary R-βHB targets, with 5 of the top 10 KEGG pathways being linked to NDDs such as AD, HD, PD, and ALS (Fig. 6g). Furthermore, all top 10 significantly enriched protein domains from primary targets of R-βHB are tubulin-related, in addition to containing the 8 tubulin-related domains found in the 1/0 mM R-βHB and F4 downregulated treatment groups (Fig. 6h).

These data together underscore the connection between our identified *ex vivo* and *in vivo* protein targets. It is evident that NDD-related proteins containing tubulin-related domains are the primary target of R-βHB, and that under subchronic BH-BD treatment conditions *in vivo* these targets are both insolubilized and cleared from the most insoluble fraction in the brain. Furthermore, the larger pool of R-βHB targets identified with 5/0 mM R-βHB treatment *ex vivo* are associated with the ubiquitin-proteasome system (UPS) *in vivo* under subchronic BH-BD treatment conditions in middle insoluble fractions in the brain. These data lay out the importance of insolubility stratification induced within the brain by R-βHB. Further elucidation of this complex R-βHB insolubilization mechanism will require careful dissection of the affected insolublome, but opens the door for exciting new avenues for utilization of ketogenic therapies in aging and NDDs.

## DISCUSSION

Here, we report evidence for a direct protein-interacting molecular mechanism of R-βHB and structurally-related metabolites in proteostasis. This activity is distinct and opposite in function from covalent posttranslational modification. We identify that βHB and other structurally similar small molecule metabolites regulate protein solubility, and that R-βHB-induced insolubilization targets include NDD-related proteins while associating with protein degradation machinery pathways *ex vivo*. Additionally, we identify that R-βHB- induced insolubilization involves structural remodeling of target proteins, and can insolubilize amyloid-β_1-42_ oligomeric structures *in vitro*, as well as high-molecular weight amyloid-β structures *ex vivo* from brain lysates from a mouse model of AD. The direct interaction of R-βHB and amyloid-β_1-42_ improves cell viability and reduces toxicity in nematode models of amyloid-β toxicity. Finally, we show an enrichment of NDD-related proteins among those insolubilized and cleared from the aged mouse brain after subchronic treatment with the ketone ester BH-BD, providing mechanistic explanation for previous literature showing ketone-related clearance of NDD-related proteins in the brain.

The observed metabolite-induced insolubilization is a robust and reproducible mechanism. While many factors can affect protein solubility *in vitro*, we showed that this mechanism is not dependent on covalent protein modification, pH, or solute load. Importantly, we reproduced the *ex vivo* effect *in vivo*, using BH-BD to deliver exogenous R-βHB to the mouse brain without other physiological alterations. R-βHB insolubilization targets that we identified *ex vivo* strongly overlap with targets found *in vivo*, supporting the similarity of mechanism between the *ex vivo* and *in vivo* systems.

Protein aggregation is a pathological feature of NDDs, but three lines of evidence support an interpretation that metabolite-induced insolubility is ameliorative rather than pathological. First, R-βHB-induced insolubility inhibits amyloid-β cytotoxicity *in vitro* with a mouse neuronal cell line. Second, *in vivo* treatment of multiple nematode models of amyloid-β proteoxicity ameliorates their phenotypes. Third, treating mice *in vivo* with BH-BD revealed clearance, rather than increased accumulation, of the most insoluble fractions of the insolublome, consistent with prior literature.

We demonstrate that a subchronic treatment with BH-BD as short as one week is sufficient to induce a broad shift in the mouse brain insolublome, with increased insolubilization of NDD-related protein targets in middle insoluble fractions (F2 and F3), and clearance of the most insolublilized aggregates (F4). We additionally identified that proteasome-related proteins were significantly enriched among F2 and F3 proteins. This association is consistent with NDD-related protein clearance observed in F4 and may shed light on the potential mechanistic details underlying clearance. We speculate that the UPS degrades proteins in fractions 2 and 3, while the autophagy lysosomal pathway degrades the more insoluble proteins in fractions 4. Protein targets identified *ex vivo* in both treatment groups were enriched for proteasome and autophagy pathways, identifying that these protein degradation machineries are key to the proteostatic activities of βHB.

The fasting metabolic state is known to be linked to proteostasis via target of rapamycin (TOR) protein kinase complex activity. TOR is activated under conditions of nutrient or energy surplus to increase translation throughput and suppress autophagy^62^. ATP itself (but not ADP) functions as a hydrotope at physiological concentrations to maintain solubility of hydrophobic proteins that might otherwise be aggregation-prone^63^. The current data implicating fasting metabolites, including R-βHB, in inducing protein insolubility to enhance degration is consistent with a broad model of cells favoring protein synthesis and stability in times of nutrient excess, and favoring repair and turnover in times of nutrient deprivation. The ability for other structurally similar small molecule metabolites to elicit insolubilization is key to understanding proteostatic improvements under alternative metabolic states. Similarities in induced insolubilization between βHB and lactate, a key metabolite upregulated during exercise and a critical fuel for neurons, may help partially explain benefits of exercise, especially in aging and NDDs^64–66^. Each metabolite may have partially overlapping but varied affinity for different proteins, providing a mechanism to both stack and finely target the proteostatic effects in individualized translation applications.

Limitations of our approach include our focus only on the brain and NDDs as systems with clear translational relevation for manipulation of proteostasis. We also focused on βHB among the set of identified metabolites because of the wide array of experimental tools available for studying ketone body biology, the well-defined role of ketone bodies in the brain, and large dynamic range of physiological concentrations of R-βHB. However, it is highly likely that metabolite regulation of protein solubility is relevant to other, if not all, tissues. Future work can define the full range of activities of the hundereds of small molecule metabolites. Futher work is also needed to define the specific brain regions and brain cell types in which metabolite-induced insolubility and clearance is most active, important, and relevant to aging and NDDs.

These data represent a missing mechanistic puzzle piece in the known literature of pathogenic protein clearance under varying metabolic states. Ketone bodies have been linked to various mechanisms of brain aging and increased healthy longevity in mice, and other fasting metabolism mechanisms have been linked to regulation of proteostasis. Here, we connect the regulation of misfolded proteins by ketone bodies with a direct molecular mechanism. It comes as no surprise that evolutionary pressures would encourage the clearance of pathogenic proteins during ketosis to promote cellular health in organisms seeking additional substrate for ATP production. In this situation, ketone bodies are janitors of damaged proteins, chaperoning away molecular waste so organisms can operate at peak molecular fitness. This mechanism can be leveraged for therapeutic development in aging and NDDs, including via pharmacological approaches for which we provide proof of principle with BH-BD. Understanding the molecular mechanisms of metabolism is an essential aspect of the future of accessible therapeutic interventions in aging and NDDs.

---

**Extended Data Fig. 1.**
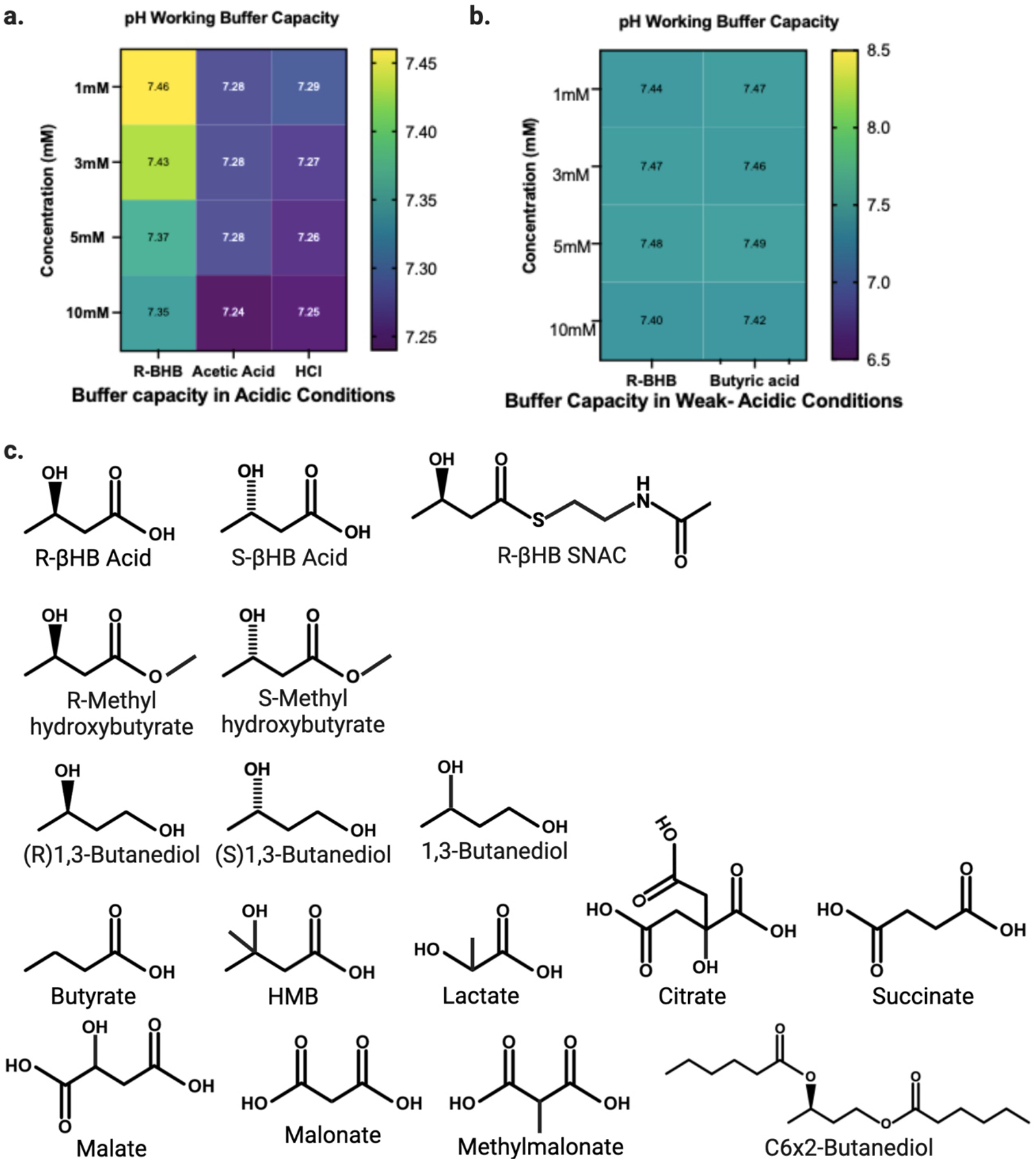
Validation of buffer capacity in experimental design. **a-b,** Quantification of pH measurements following equimolar acid addition to TEM buffer with **(a)** strong acids and **(b)** weak acids. **c,** Chemical structures of compounds used throughout study.

**Extended Data Fig. 2.**
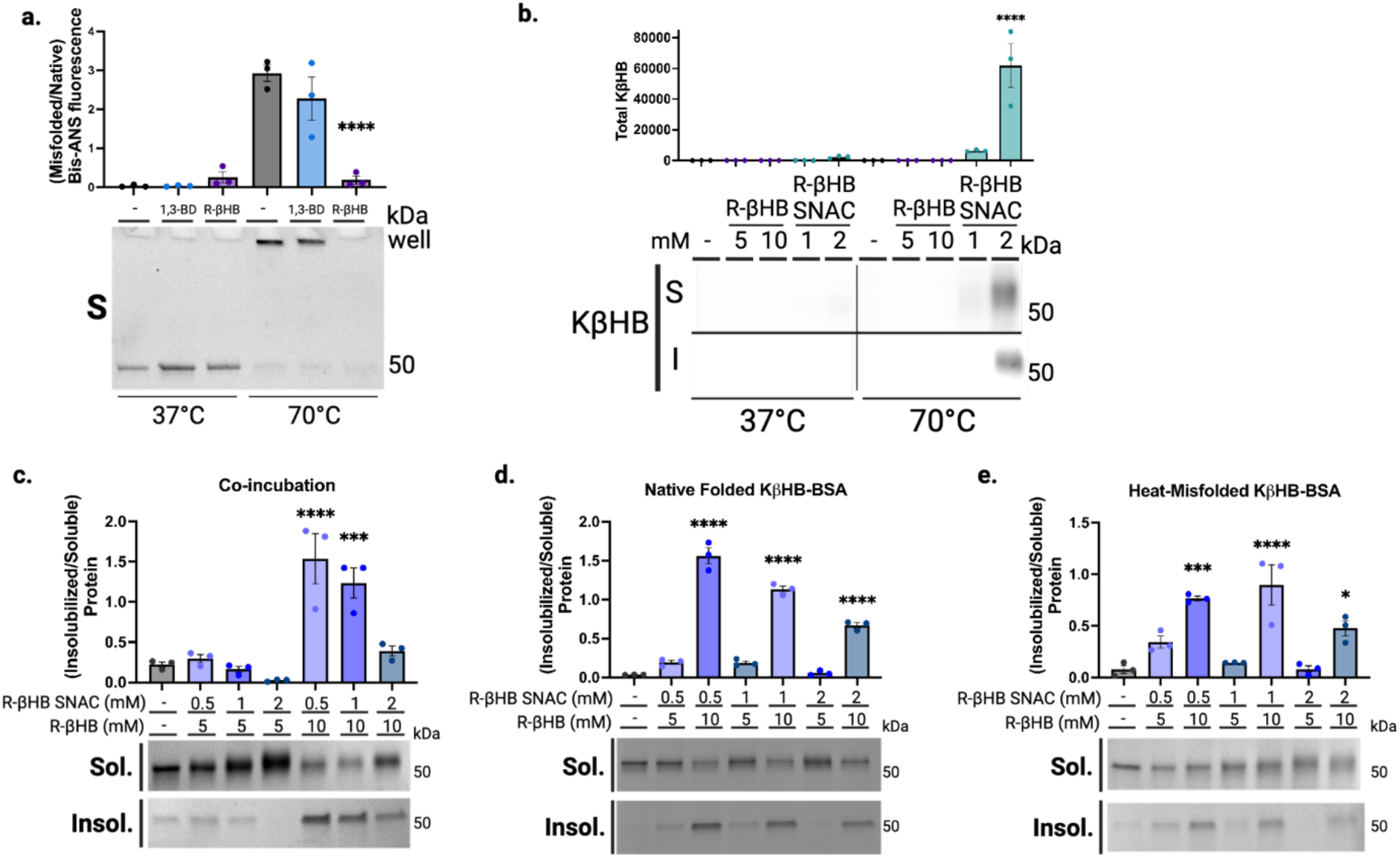
β-hydroxybutyrate directly induces protein insolubility without posttranslational modification. **a,** SDS-PAGE and quantification of Bis-ANS fluorescence in soluble fraction of native (50 kD, +37°C) and heat-misfolded (aggregated in well, +70°C) BSA treated with 10 mM of R-βHB and 10 mM of 1,3-BD. **b,** Western blot and quantification for KβHB in native (+37°C) and heat-misfolded (+70°C) BSA which remains soluble or becomes insolubilized following exposure to 5-10 mM of R-βHB and 1-2 mM of R-βHB-SNAC. **c-e,** Imperial staining and quantification of BSA which remains soluble or is insolubilized following **(c)** simultaneous exposure of BSA to 5-10 mM of R-βHB and 0.5-2 mM R-βHB-SNAC at +70°C, **(d)** native folded (+37°C) KβHB-BSA treated with 5-10 mM of R-βHB, and **(e)** heat-misfolded (+70°C) KβHB-BSA treated with 5-10 mM of R-βHB. a, Representative image from triplicate repetitions. b, Mean ± S.E.M, N=3, p-value calculated using one-way ANOVA with Sidak’s multiple comparison test. c, Mean ± S.E.M, N=3, p-value calculated using one-way ANOVA with Dunnett’s multiple comparison test. d, Mean ± S.E.M, N=3, p-value calculated using one-way ANOVA with Dunnett’s multiple comparison test. e, Mean ± S.E.M, N=3, p-value calculated using one-way ANOVA with Dunnett’s multiple comparison test.

**Extended Data Fig. 3.**
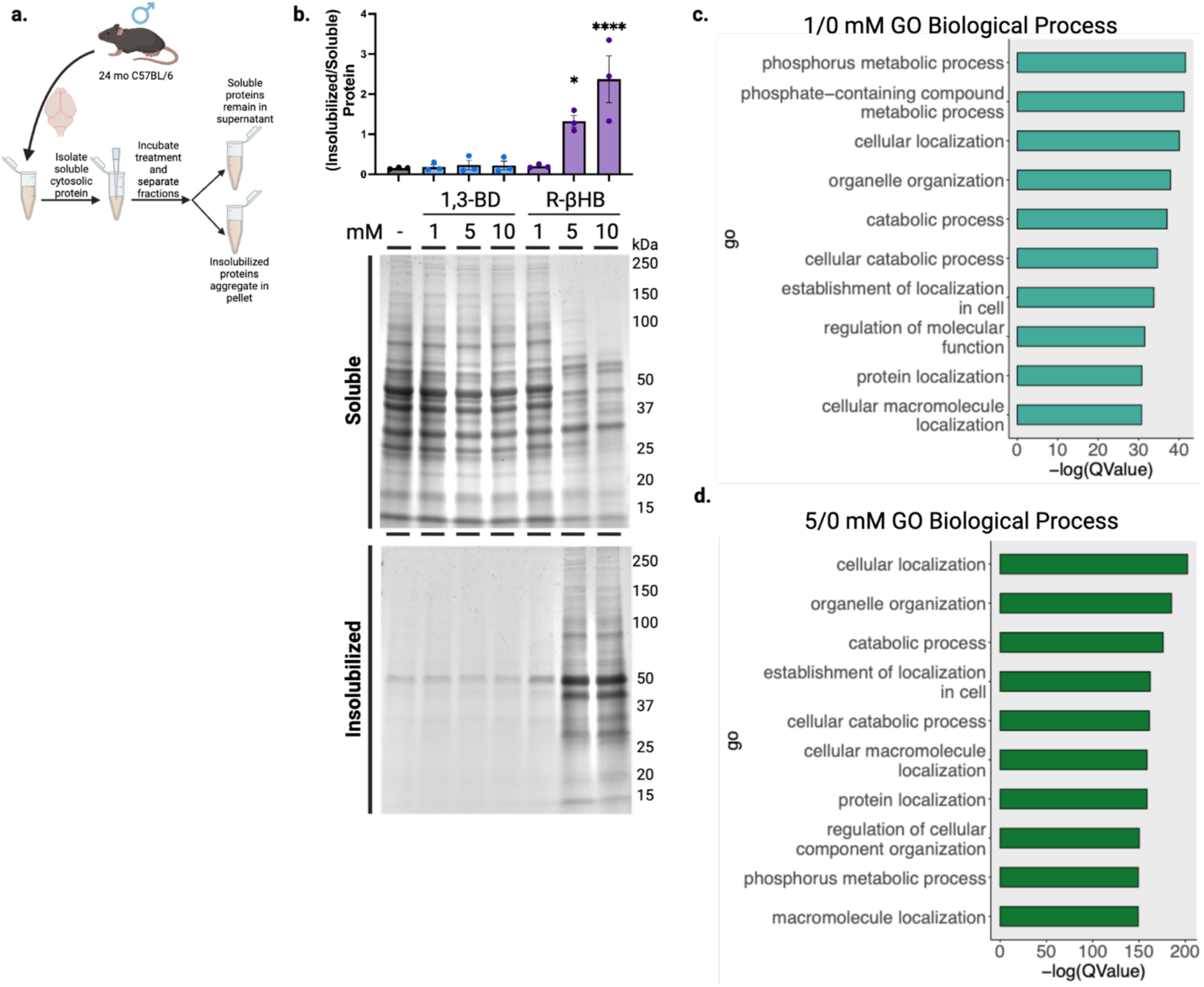
β-hydroxybutyrate remodels the older C57BL/6 mouse brain proteome *ex vivo* via insolubilization of targets. **a,** Schematic of *ex vivo* protein insolubilization assay. **b,** Imperial staining and quantification of 24 month female wild-type mouse soluble cytosolic brain proteins which remain soluble or are insolubilized after treatment with 1-10 mM of R-βHB and 1-10 mM of 1,3-BD. **c-d,** Dotplots from clusterProfiler gene ontology overrepresentation analysis for biological process on top 10 significantly upregulated terms in **(c)** 1/0 mM and **(d)** 5/0 mM R-βHB treatment groups. b, Mean ± S.E.M, N=3, p-value calculated using one-way ANOVA with Dunnett’s multiple comparison test.

**Extended Data Fig. 4.**
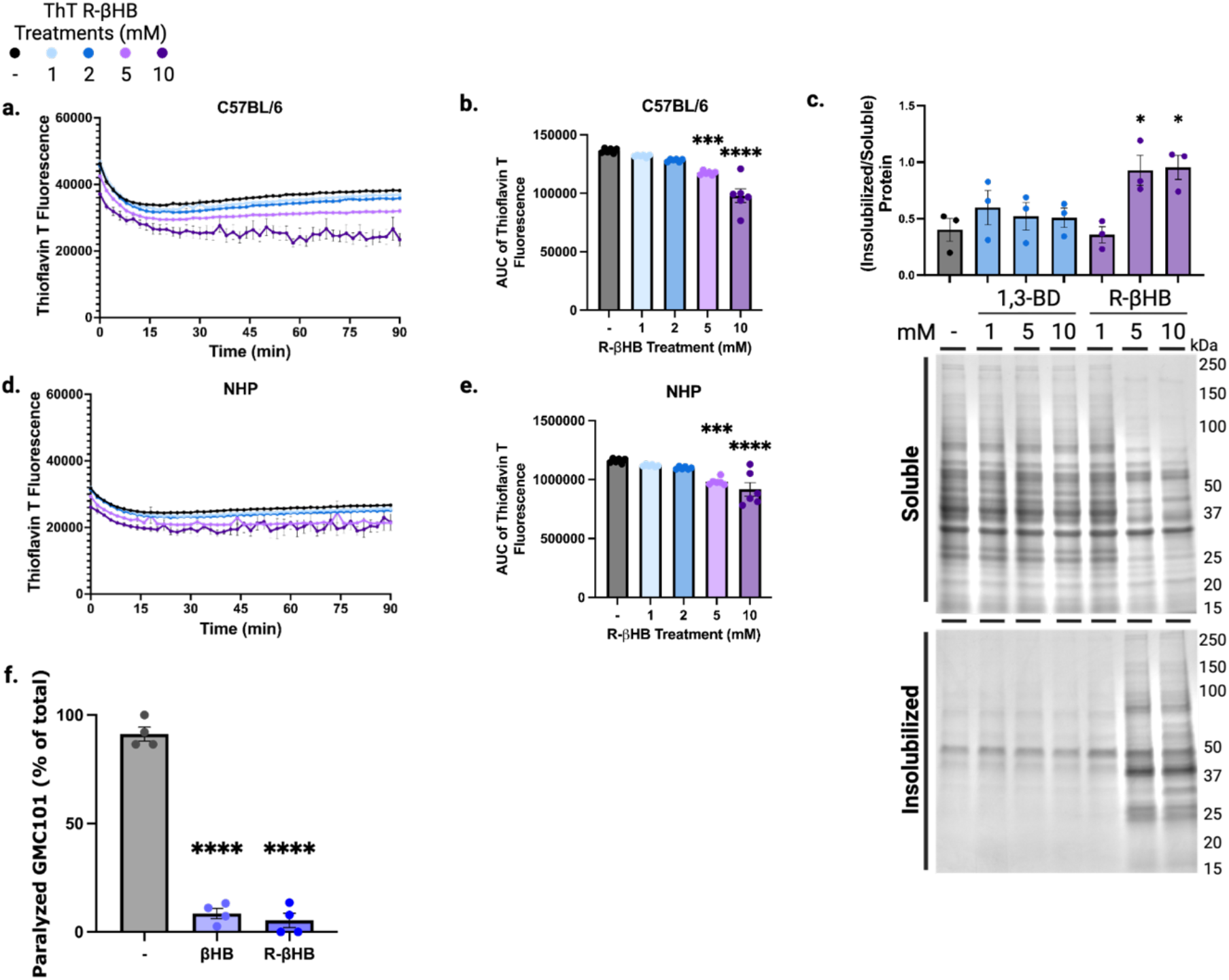
β-hydroxybutyrate inhibits oligomeric toxicity through structural remodeling of proteins and suppresses human amyloid-β-induced paralysis and neurotoxicity in *C. elegans*. **a-b,** Total protein aggregation kinetics monitored via Thioflavin T fluorescence in mouse soluble cytosolic brain proteins treated with 1-10 mM of R-βHB from 24 month wild-type female, **(a)** timecourse and **(b)** area under the curve. **c,** Imperial staining and quantification of 25 year female non-human primate soluble cytosolic brain proteins which remain soluble or are insolubilized after treatment with 1-10 mM of R-βHB and 1-10 mM of 1,3-BD. **d-e,** Total protein aggregation kinetics monitored via Thioflavin T fluorescence in non-human primate soluble cytosolic brain proteins treated with 1-10 mM of R-βHB from 25 year rhesus macaque female, **(d)** timecourse and **(e)** area under the curve. **f,** Quantification of amyloid-β proteotoxicity in temperature-sensitive (aggregation-permissive at +25°C) GMC101 strain, determined by scoring the percentage of animals paralyzed at 33-34 hours following temperature shift and with 50 mM of βHB treatment, all bacteria was UV treated. a, Mean ± S.E.M, N=6, p-value calculated using one-way ANOVA with Dunnett’s multiple comparison test. b, Mean ± S.E.M, N=6, p-value calculated using one-way ANOVA with Dunnett’s multiple comparison test. c, Mean ± S.E.M, N=3, p-value calculated using one-way ANOVA with Dunnett’s multiple comparison test. d, Mean ± S.E.M, N=6, p-value calculated using one-way ANOVA with Dunnett’s multiple comparison test. e, Mean ± S.E.M, N=6, p-value calculated using one-way ANOVA with Dunnett’s multiple comparison test. f, Mean ± S.E.M, N=4 (∼300 animals). p-value calculated using one-way ANOVA with Dunnett’s multiple comparison test.

**Extended Data Fig. 5.**
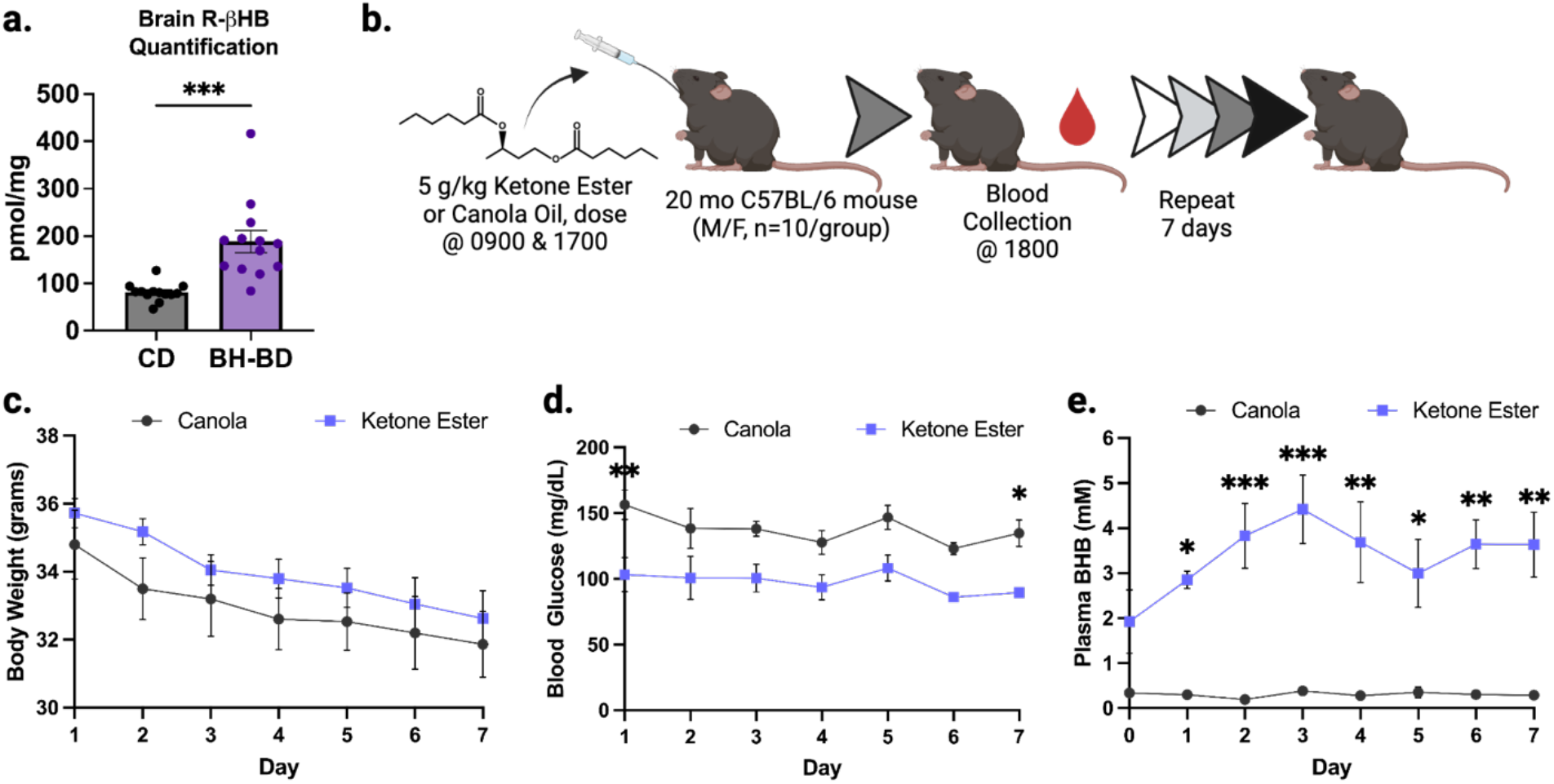
Subchronic treatment with a ketone ester induces ketosis. **a,** Absolute quantification of R-βHB in brain tissue of 22 month male and female wild-type mice fed for 7 days with control diet or BH-BD. **b,** Schematic of subchronic BH-BD and control treatment schedule. **c,** Quantification of animal body weight at 0800 daily. **d,** Quantification of blood glucose concentrations 1 hour post oral gavage. **e,** Quantification of plasma βHB concentrations 1 hour post oral gavage. a, Mean ± S.E.M, N=13, p-value calculated using Kolmogorov-Smirnov test. c, Mean ± S.E.M, N=5-6, p-value calculated using mixed-effects analysis with Sidak’s multiple comparison test. d, Mean ± S.E.M, N=5-6, p-value calculated using mixed-effects analysis with Sidak’s multiple comparison test. e, Mean ± S.E.M, N=5-6, p-value calculated using mixed-effects analysis with Sidak’s multiple comparison test.

**Extended Data Fig. 6.**
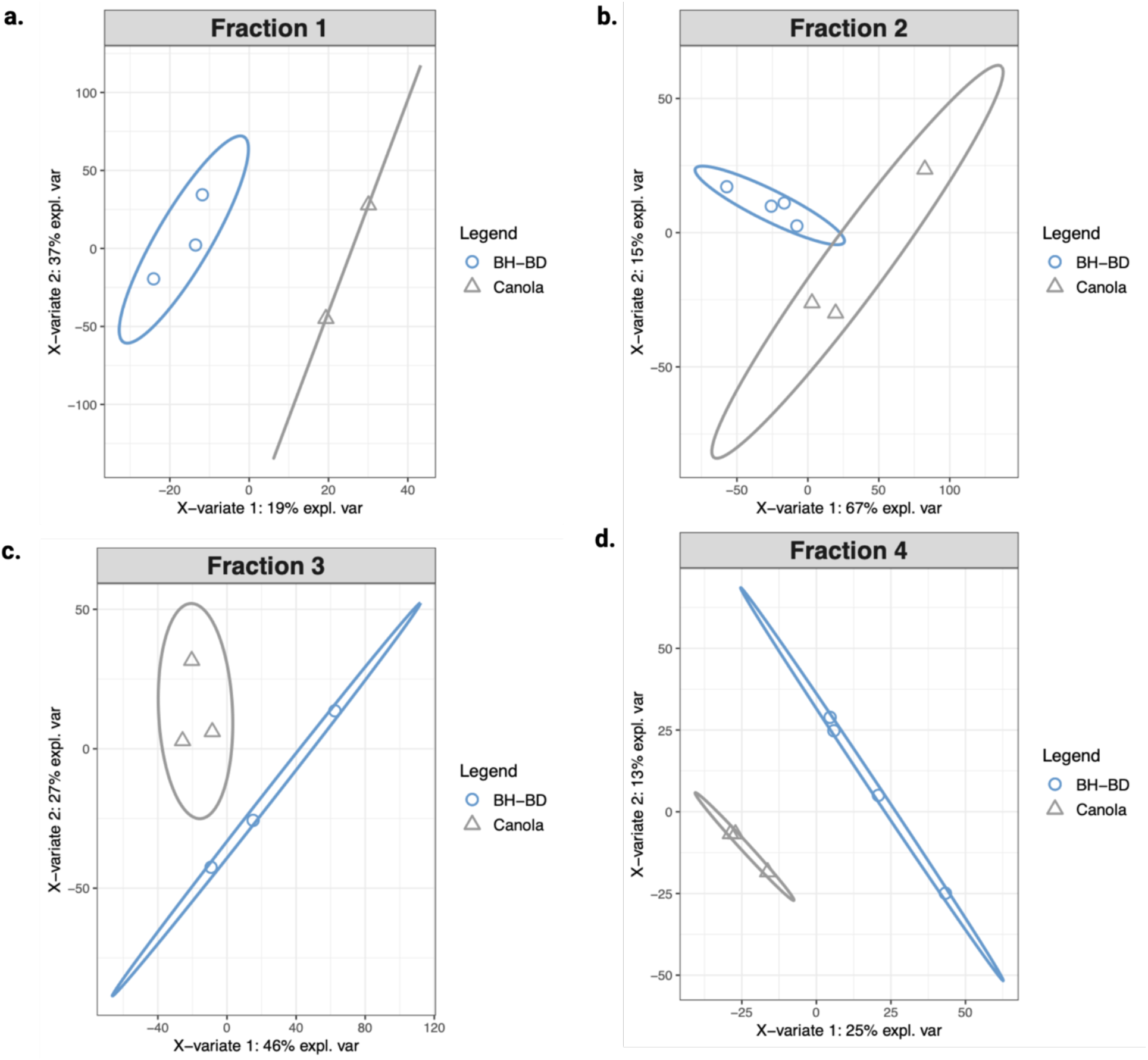
Subchronic treatment with BH-BD restructures the insolublome. **a-d,** Partial least squares-discriminant analysis (PLS-DA) of *in vivo* BH-BD/control proteomic samples, **(a)** Fraction 1, **(b)** Fraction 2, **(c)** Fraction 3, and **(d)** Fraction 4.

**Extended Data Fig. 7.**
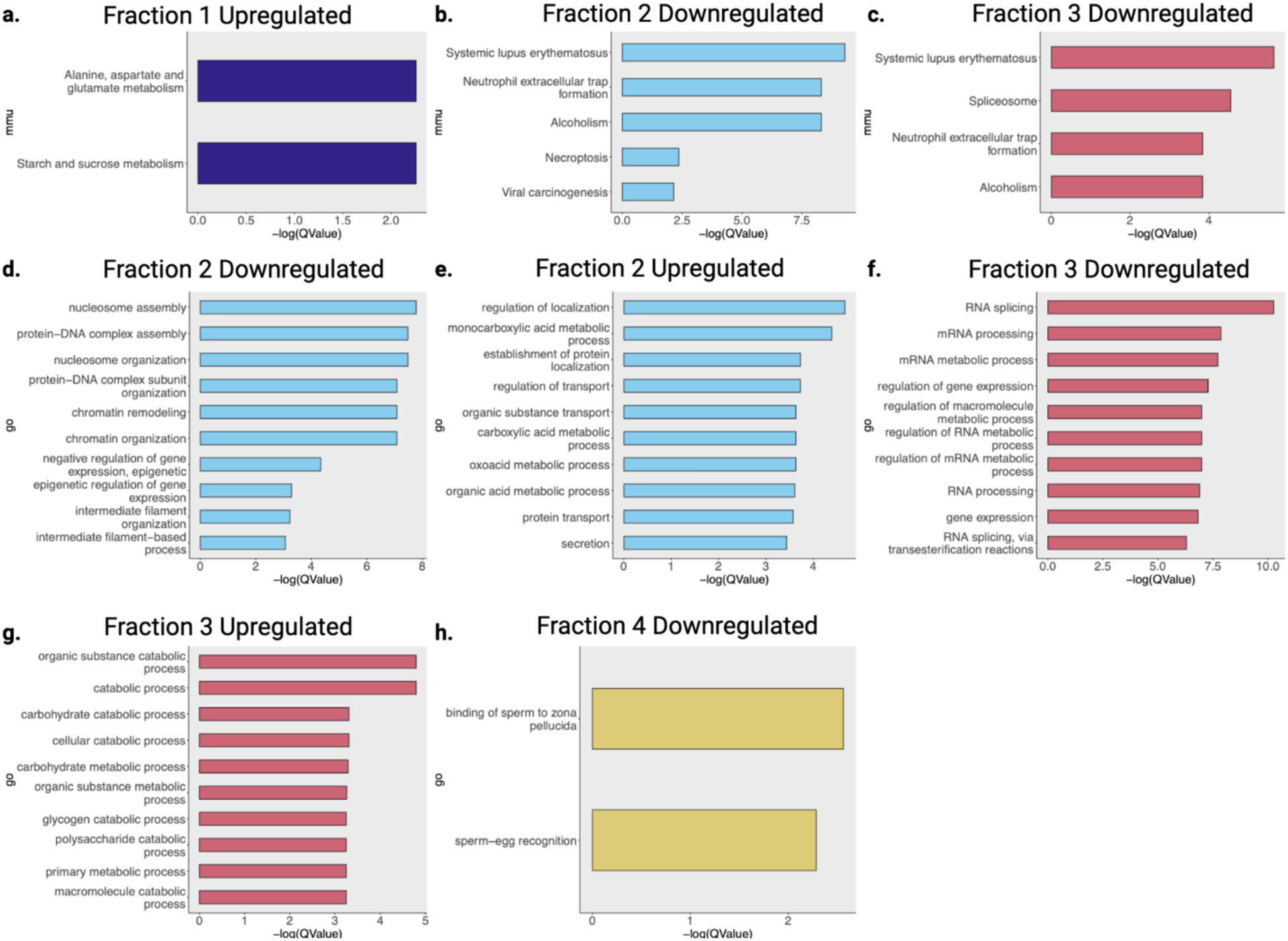
Subchronic treatment with a ketone ester elicits changes aged mouse brain insolublome. **a-c,** Dotplots from clusterProfiler KEGG pathway overrepresentation analysis on significantly **(a)** upregulated proteins from Fraction 1, **(b)** downregulated proteins from Fraction 2, and **(c)** downregulated proteins from Fraction 3. **d-h,** Dotplots from clusterProfiler gene ontology overrepresentation analysis biological process terms from **(d)** downregulated proteins in Fraction 2, **(e)** upregulated proteins in Fraction 2, **(f)** downregulated proteins in Fraction 3, **(g)** upregulated proteins in Fraction 3, and **(h)** downregulated proteins in Fraction 4.

## METHODS

### Figure panels

Figure panels were developed in R Studio (Version 4.2.2, 2022_10_31, “Innocent and Trusting”) or GraphPad Prism (Version 8.2.1(279) and 9.4.1(458)) and were imported into BioRender for final formatting.

### Mice

C57BL/6 mice were acquired from the National Institute of Aging’s Aged Rodent Colony. Breeding pairs of hAPPJ20^67^ mice were provided from Jorge Palop (Gladstone Institutes, San Francisco, USA). Mice were allowed to acclimate to vivarium environmental conditions for at least 2 weeks prior to use in any experiment. Mice were randomly assigned to groups at the beginning of each experiment. Aged wild-type mice were 20-24 months and J20 mice were 4 months. Mice were housed at +22.2°C and 52.1% humidity. Under a 12-hour light-dark cycle, mice were kept in filter-topped cages with autoclaved food and water at the Buck Institute for Research on Aging in Novato, California. All experiments were performed in accordance with guidelines set by facilities and were approved by the regulations of the institutional animal care and use committee (IACUC).

### Mouse tissue collection

Mice were euthanized by CO_2_, followed by bilateral thoracotomy, and tissues were immediately collected, sub-dissected as needed, and flash-frozen via liquid nitrogen in 2.0 mL cryogenic vials. Vials were then transferred to −80°C for long term storage.

### Non-human Primates

Rhesus macaque brain tissue was received as fresh-frozen tissue blocks from the National Institute of Aging Non-human Primate Tissue Bank (NIA NHP). Further handling was at Biosafety Level 2 to prevent transmission of pathogens and storage was at −80°C.

### Tissue homogenization and subcellular fractionation

Pre-sectioned, frozen wild-type mouse, J20 mouse, or non-human primate brain tissue was weighed and immediately homogenized in a 2.0 mL glass mortar and pestle (Corning 7727-02 Pyrex) with a ratio of 1 mg tissue to 5 µL cold TEM buffer (50 mM Tris, 1 mM EDTA, 0.5 mM MgCl_2_, pH 7.4) + 1x protease inhibitor cocktail (Abcam ab271306) with 30 up-and-down strokes on ice. Homogenate was transferred by pipetting into a 1.5 mL microtube and centrifuged at 2,000xg for 20 minutes at +4°C. The pellet (P1) was discarded. The supernatant (S1) was transferred by pipetting to a polypropylene tube (Beckman Coulter 326819) and centrifuged at 100,000xg for 60 minutes at +4°C. The resulting supernatant (S2) was transferred by pipetting to a 1.5 mL microtube and stored at −80°C until further usage. The pellet (P2) was resuspended by up-and-down pipetting in an equal volume of NDSD buffer (50 mM Tris, 1 mM EDTA, 0.5 mM MgCl_2_, 0.5% NP40, 0.5% Sodium Deoxycholate, 1% SDS, 1 mM DTT), followed by incubation overnight at +25°C and centrifugation the next day at 20,000xg for 20 minutes at +25°C to produce the final resolubilized proteins in the supernatant, stored afterwards at −80°C until further usage.

### *In vitro* bovine serum albumin insolubilization assays

Bovine serum albumin (BSA) standard ampules (Pierce 23209) were opened and BSA was pipetted to a 1.5 mL microtube where dilution to 1.0 mg/mL was performed with milli-Q water. 1.0 mg/mL BSA was combined with fresh TEM buffer (without protease inhibitor cocktail) and master stock of metabolite treatment compounds to achieve final volume of 50 µL in a 1.5 mL microtube. BSA was added after metabolite treatment was diluted in TEM buffer to prevent exaggerated or local binding effects (first pipetting metabolite, then TEM, then BSA). Microtubes were briefly vortexed and spun to collect volume at bottom of tube prior to incubation for 60 minutes at noted temperatures in a thermomixer (Eppendorf ThermoMixer C). Microtubes were immediately transferred and centrifuged at 20,000xg for 60 minutes at +25°C. All supernatants were transferred to a fresh 1.5 mL microtube, and all pellets were resuspended in an equal volume of NDSD buffer (without DTT) as described in *Tissue homogenization*. All samples were stored at −80°C until further usage. In some experiments after incubation, BSA samples were transferred to Amicon ultra-0.5 mL centrifugal filter units with 3 kDa molecular weight cutoff (Millipore Sigma UFC5003) and filtered according to manufacturer instructions to collect posttranslationally modified BSA for secondary incubation.

### *Ex vivo* protein insolubilization assay

Post-ultracentrifugation supernatant (S2) from brain lysates was thawed on ice and combined with fresh TEM buffer + 1x protease inhibitor cocktail and 100 mM stock of metabolite treatment compounds to achieve final volume of 50 µL in a 1.5 mL microtube. S2 was added after metabolite treatment was diluted in TEM buffer as to prevent exaggerated or local binding effects (first pipetting metabolite, then TEM, then TEM-buffered S2). 20 µL (∼4 mg/mL) of supernatant was used to achieve a final concentration of ∼1.6 mg/mL per reaction. Microtubes were briefly vortexed and spun to collect volume at bottom of tube prior to incubation at 300 rpm for 60 minutes at +37°C in a thermomixer. Microtubes were immediately transferred and centrifuged at 20,000xg for 60 minutes at +25°C. All supernatants were transferred to a fresh 1.5 mL microtube, and all pellets were resuspended in an equal volume of NDSD buffer as described in *Tissue homogenization*. All samples were stored at −80°C until further usage.

### Gel electrophoresis, protein staining, bis-ANS fluorescence, and immunoblotting

Gel electrophoresis was performed using SDS-PAGE. Post-assay, isovolumetric loading samples were prepared by combining assay samples with 4x LDS sample buffer (Invitrogen NP0007) + β-mercaptoethanol (9:1, LDS:βME). Isovolumetric assay samples were used, instead of normalizing by protein concentration, as it is critical to maintain consistency from input. During previous assay, all reactions were normalized by protein quantity, differences in insolubilization are expected based on treatment and unequal concentrations are therefore meaningful. When using samples from lysate or other assays, protein concentration was quantified using bicinchoninic acid (BCA) protein assay (Pierce 23227) and ≥10 µg protein was loaded per well. After combining with LDS + βME buffer, loading samples were vortexed and spun down, boiled at +75°C for 10 min, vortexed and spun down again, then finally loaded into BIORAD Criterion TGX Precast gels (4-20%). 2 µL of protein ladder (BIORAD 1610374) was used. Gels were run in a BIORAD Criterion cell (mini and midi format) with running buffer for 10 min at 80 V and 60 min at 120 V, or until dye front has nearly run off, using a power supply.

Protein staining was performed immediately after completion of gel electrophoresis. Gels were cut from plate and immediately placed in Imperial stain (Thermo 24615) for 60 min at +25°C or microwaved repeatedly for 15 s until fully stained. Gels were then de-stained in milli-Q water overnight and imaged.

4,4’-dianilino-1,1’-binaphthyl-5,5’-disulfonic acid (bis-ANS) fluorescence was measured after completion of gel electrophoresis using an adapted protocol from previous literature^57^. Samples from *in vitro* bovine serum albumin assay were incubated with 500 µM bis-ANS (100 µM final) in a black 96 well plate under a UV transilluminator torch (standard filter), fully covered from external light. After activation, samples were loaded into the BIORAD Criterion TGX Precast gels (4-20%) as described and run in darkness. After completion of gel electrophoresis, fluorescence was imaged at 488 nm.

Immunoblotting was performed immediately after completion of gel electrophoresis. Gels were trimmed and sandwiched against 0.2 µm nitrocellulose blotting membrane (Prometheus 84-875) and blot paper (WypAll X60). Blotting membrane was primed for 3 min prior in cold transfer buffer (BIORAD 10026938), according to manufacturer specifications. Blot paper was soaked in cold transfer buffer immediately before sandwiching. Sandwich was rolled to remove bubbles before closing of cassette. BIORAD TransBlot Turbo transfer system was used for all transfers. After transfer, membranes were cut to remove excess and Ponceau S stained (VWR K793) to confirm complete transfer. Once completely de-stained in 0.1 N NaOH, membranes were housed in clear blotting boxes and blocked in 5% blocking buffer in TBS-T (BIORAD 1706404) for 40 min at +25°C. After completion of blocking, membranes were incubated in primary antibody diluted in 5% blocking buffer overnight for 16 hours at +4°C with continuous gentle rocking. Primary antibodies used targeted pan anti-β-hydroxybutyryllysine (PTM Biolabs, PTM-1201) or anti-amyloid-β_1-16_ (BioLegend 6E10, 803004). After primary antibody incubation, antibody dilutions were saved at −20°C for up to 5 re-uses. Membranes were next washed in TBS-T 3 times for 10 min each before blocking in 1:2000 secondary antibody for 60 min at +25°C. Membranes were again washed in TBS-T 3 times for 10 min each, or more until TBS-T is visually clear, before imaging. Imaging was performed with enhanced chemiluminescent detection (Thermo 34096) on Azure Biosystems c600 imager.

Stain and immunoblot images were exported to Image Studio and corrected for contrast to show appropriately similar banding and background for easier quantification. Images were next transferred to ImageJ and bands were quantified using densitometry, with correction normalization by background or control in Microsoft Excel. Values were finally exported to GraphPad Prism for plotting.

### *In vitro* protein conformation assay

Post-ultracentrifugation supernatant (S2) from brain lysates were thawed on ice alongside 1 mM Thioflavin T (VWR 103802-652). Thioflavin T (ThT) was prepared by resolubilizing powder in quickly moving milli-Q water, then aliquoted into foil-wrapped 1.5 mL microtubes and stored at −20°C until further use. Master-mix was created using the following volumes per well: 33 µL fresh TEM buffer + 1x protease inhibitor cocktail was mixed with 10 µL post-ultracentrifugation supernatant on ice. 2 µL ThT per well was added to master-mix just before pipetting into the well to prevent exaggerated or local binding effects that were not representative of the assay. Metabolite treatment compounds were added directly to well with necessary TEM buffer to bring volume to 5 µL per well. Master-mixes were all added to one well and reverse-pipetted thrice into the three replicate wells; master-mix well readings were not used for data analysis. Final reaction volume in each well was 50 µL. All assays were run using Corning 3904 plates and imaged every 2 min in a CLARIOstar Plus microplate reader with an excitation wavelength of 444 and an emission wavelength of 491 with double-orbital shaking before reading and between reading at 300 rpm. Timepoint values were exported to Microsoft Excel and transposed to GraphPad Prism for plotting.

### Amyloid-β peptide incubation in aged wild-type brain environment

0.1 mg of lyophilized HiLyte Fluor 488-labeled amyloid-β_1-42_ (Anaspec AS-60479-01) was dissolved in 50 µL of 1% NH_4_OH and diluted to 0.5 mg/mL with 1xPBS before aliquoting for storage at −20°C until further use. Aliquots were thawed on ice and combined with TEM buffer + 1x protease inhibitor cocktail to dilute amyloid-β_1-42_ to a final concentration of 0.33 mg/mL. Compounds were added at specified concentrations, post-ultracentrifugation supernatant (S2) from wild-type mouse brain lysate was added to reach a final concentration of ∼1.6 mg/mL per reaction, and solution was incubated at +37°C for 60 minutes in a thermomixer at 300 rpm before centrifugation at 25,000xg for 60 minutes at +25°C. All supernatants were transferred to a fresh 1.5 mL microtube, and all pellets were resuspended in an equal volume of NDSD buffer (without DTT) as described in *Tissue homogenization*. All samples were stored at −80°C until further usage.

### Amyloid-β oligomer generation

To make 200 μM of amyloid-β monomer stock, 1 mg of lyophilized amyloid-β_1-42_ (Cayman 20574) was dissolved in 40 μL of 1% NH_4_OH and 500 μL of 1xPBS, then stored at −20°C until further use. To produce amyloid-β oligomers, the monomer stock was incubated with 50 μL of TEM Buffer, phosphatase inhibitor (PI), and 30 μL of incubation media (DMEM (Gibco 11966025), 1 mM glucose, penicillin/streptomycin (Corning 30-002-CI)) for 2 hours at +37°C in darkness.

### Cell culture

N2a cells (ATCC CCL-131) were maintained in culture media (DMEM (Corning 10-013-CV), 10% FBS (Corning 35-011-CV), and penicillin/streptomycin) in a +37°C incubator with 5% CO_2_. The cells were then cultured in a 96-well plate (30,000 cells/well) with culture media. On the next day, the cells were incubated with and without incubation media, TEM + PI, amyloid-β oligomers (final concentration of 2 μM), and R-βHB (Sigma 54920) (final concentration of 10 mM) for 24 hours in the incubator. The cell proliferation was quantified using a XTT Assay Kit (Cayman 10010200) following the manufacturer protocol.

### *C. elegans* strain information

N2 - wild type, GMC101 - dvIs100 [unc-54p:: amyloid-β_1-42_::unc-54 3’-UTR + mtl- 2p::GFP], UA198 - *baIn34*[P*_eat-4_*::Aβ,P*_myo-2_*::mCherry]; *adIs1240*[P*_eat-4_*::GFP]

*C. elegans* strains were maintained at +20°C under standard laboratory conditions as described previously^68^. For experimental purposes worms were developmentally synchronized from an egg lay of 3 hours. Please see the figure legends for details of trials and statistics used to determine significance.

### *C. elegans* preparation of plates

A 2 M stock of βHB sodium (Acros Organics) was aliquoted and stored at −20°C. 100 µL of working solution (50 mM βHB) was prepared by mixing 75 µL of stock with 25 µL of sterile water and was added to the top of the 35 mm NGM plates (3 mL NGM agar) already seeded with a bacterial OP50 lawn. For control plates, 100 µL of sterile water was added to the top of the 35 mm NGM plates (3 mL NGM agar). Experiments with heat killed bacteria utilized liquid OP50 bacterial culture which was incubated at +70°C for 60 minutes with occasional shaking to seed plates. We chose to add compounds on top of the seeded plates and not in the agar plate to ensure maximum bioavailability and ensure stability of the compounds. The bacterial feeding solution is akin to food rather than an intravascular or intracellular space, and compounds are diluted upon ingestion and circulation in the animals. Therefore, higher βHB concentrations are used on the plate than in cell culture media or measured in mammalian blood. For example, maximal extension of *C. elegans* lifespan is at 20 mM βHB in the feeding solution^69^. Plates were allowed to sit at +20°C for 24 hours before use or before moving into +4°C. Plates were stored for no longer than one week.

### *C. elegans* paralysis assay

Egg-lay synchronized populations of GMC101 (expresses human amyloid-β_1-42_ protein in the body wall muscles^70^ were grown from eggs at +20°C. 68-72 hours after egg-lay, animals were transferred to fresh 35 mm plates treated with control (water) or 50 mM βHB. Plates were immediately shifted to +25°C and paralysis was scored 24-28 hours after the temperature shift when control animals reached 90% paralysis. Animals were scored as paralyzed if they failed to move either spontaneously or if they failed to respond to touch-provoked movement with a platinum wire.

### *C. elegans* glutamatergic neurodegeneration assay

Egg-lay synchronized populations of UA198 (expresses human amyloid-β_1-42_ and GFP in glutamatergic neurons^71^ were grown from eggs at +20°C. 68-72 hours after egg-lay, day-1 young adult animals were transferred to control (water) or 50 mM βHB plates. The young adult UA198 strain when visualized under fluorescent microscope shows GFP expression that marks 5 intact glutamatergic neurons in their tail-region. Expression of amyloid-β_1-42_ results in age-dependent neurodegeneration. Animals were scored for the presence of 5 intact glutamatergic neurons at day-3 and day-5.

### Mouse diets, feeding, and BH-BD oral gavage

All mice were given access to food (diets from Envigo) and water ad libitum and were only supplemented with additional compounds as noted. Chow diet (Teklad 2918, Irradiated Global 18% Protein Rodent Diet) was sourced from the vivarium at Buck Institute for Research on Aging and was unmodified. Ketone ester used was bis-hexanoyl (R)-1,3-butanediol (BH-BD), supplemented in noted concentrations per experiment. Teklad TD.150345 (93M, Irradiated) was used as a control diet (CD) for comparison to fed BH-BD. BH-BD was synthesized by WuXi Apptec (China) at >98% purity and has a light-yellow appearance. Rodent metabolism, kinetics, and safety data have previously been reported^60, 72^. For gavage, a body weight adjusted amount of undiluted BH-BD was loaded into syringes and administered via oral gavage using 20 g, 1.5 in curved, 2.25 mm ball reusable stainless feeding needles (Braintree Scientific) at times 0900 and 1700. Gavage control groups were administered a volume matched canola oil dose (Wesson) to ensure isocaloric treatment between groups. Mice were awake and under no anesthetic for oral gavage administration. Body weight was measured prior to 0900 and was used to calculate oral gavage volume for individual mice. Chow food weight was measured prior to 0900 and was noted to identify weight loss in groups. Euthanasia was immediately after final blood collection at 1800, one hour after final gavage on the 7^th^ day of timecourse.

### Mouse diets, feeding, and absolute quantification of brain (R)-βHB

Animals from this experiment were fed with either CD or BH-BD, food was provided ad libitum at all times. Per-calorie macronutrient content for customized diets (Envigo) is as follows: CD, 77% carbohydrate, 13% fat, 10% protein (TD.150345); BH-BD (31% w/v BH-BD), 53% carbohydrates, 9% fat, 7% protein. All mice were acclimated on CD for 2 weeks in the groups they arrived before they were single-caged for this study. Mice in this study are male (N=7) and female (N=6). Frozen whole brains (∼30-50 mg) were homogenized in extraction buffer [3:1, v/v acetonitrile and HPLC-grade H2O] with Next Advance Bullet Blender (BBY24M). Derivatization was adapted from Tsutsui et al^73^. Extracted samples were dried using DNA SpeedVac System (ThermoFisher Scientific Model DNA130-115) and resuspended in 98:2 H2O:Methanol containing 0.1% formic acid, mixed and centrifuged at 10,000xg for 10 minutes. Supernatant was then transferred into HPLC vials. A sample volume of 2 μL was injected into the UPLC-MS/MS Thermo Q Exactive with Vanquish Horizon in Full MS and PRM scan modes using positive ionization. The analysis was performed on a Accucore Vanquish C18+ column (100 x 2.1 mm, 1.5µm particle size; ThermoFisher Cat. #20073385). The following mobile phases were used: A) HPLC-graded H2O containing 0.1% (v/v) formic acid and B) methanol containing 0.1% (v/v) formic acid with the gradient starting at 2% B for 0.5 min and gradually increasing to 10.4%B until 10.0 min, 2%B until 10.1 min, and 2%B until 12.0 at 0.150 mL/min flow rate for a total run time of 12.0 minutes. Column was maintained at +40°C. LC system was hyphenated to Thermo Q Exactive MS equipped with heated electrospray ionization (HESI) source. The MS system was operated in Full Scan MS or PRM modes using positive ionization. MS scan range was 50.0 to 750.0 m/z in Full MS scan mode. The resolution was set to 140,000 with AGC target 3e6, isolation window 1.0 m/z and optimal collision energy was 50 (arbitrary units). For PRM scan mode the isolation window was set to 0.4 m/z and resolution was set to 70,000 with AGC target 1e6. Common HESI parameters were auxiliary gas: 5, sheath gas flow: 50, sweep gas: 0, spray voltage 3 kV, capillary temperature +320°C, S-lens 55.0, and auxiliary gas temperature: +150°C. Quantification of area response ratios were processed and acquired using Thermo Scientific Xcalibur software (OPTON-30965). Area Response Ratios (ES/IS) from three technical replicates were averaged, and a simple regression line was constructed. To quantify the ’samples’, Area Response Ratios (ES/IS) from three technical replicates were averaged and concentrations were calculated using the Prism-generated calibration curve equations. Amounts were calculated as [(100*concentration) pmol]/[mg tissue weight].

### Mouse metabolic data collection

Blood was obtained via distal tail-snip and immediately used for glucose measurements; additional blood was collected in lithium-heparin coated microvettes (Sarstedt CB 300 LH) and kept on ice. Afterwards, samples were centrifuged at 1,500xg for 5 min at +4°C to separate plasma, which was kept at −80°C until further usage. Previous experiments have confirmed no freeze-thaw interference effects. βHB plasma concentrations were measured using a colorimetric, benchtop assay (Stanbio 2440058), using 3 µL plasma volumes in triplicate.

### Sequential Detergent Extraction of Aggregates

Protocol was adapted from Shaw et al^74^. Pre-sectioned, frozen wild-type mouse brain tissue from the BH-BD oral gavage cohort was weighed, immediately homogenized, and differentially centrifuged as described in *Tissue homogenization and subcellular fractionation*. After ultracentrifugation at 100,000xg for 60 minutes at +4°C, the pellet (P2) was instead resuspended in TEM + 0.5% NP40. The resolubilized solution was incubated at +37°C for 30 minutes in a thermomixer before centrifugation at 25,000xg for 30 minutes at +25°C, the supernatant is fraction 1 (F1) and the pellet (P3) was resuspended in TEM + 0.5% NP40 + 0.5% Sodium Deoxycholate + 0.25% SDS. The resolubilized solution was incubated at +37°C for 30 minutes in a thermomixer before centrifugation at 25,000xg for 30 minutes at +25°C, the supernatant is fraction 2 (F2) and the pellet (P4) was resuspended in TEM + 0.5% NP40 + 0.5% Sodium Deoxycholate + 2% SDS. The resolubilized solution was incubated at +37°C for 30 minutes in a thermomixer before centrifugation at 25,000xg for 30 minutes at +25°C, the supernatant is fraction 3 (F3) and the pellet (P5) was resuspended in TEM + 0.5% NP40 + 0.5% Sodium Deoxycholate + 3% SDS. The resolubilized solution was incubated at +25°C overnight before centrifugation at 25,000xg for 30 minutes at +25°C, the supernatant is fraction 4 (F4).

### Protein digestion and desalting

Aliquots of sequential detergent extraction fractions (F1-F4) varying from 2 to 24.5 µg were brought up to the same overall volume of 50 µL with water. Cell pellets from Ex Vivo protein insolubilization assay were resuspended in 100 µL of 0.5% SDS in 100 mM triethylammonium bicarbonate buffer (TEAB) with 1x protease inhibitor cocktail, using ∼10 µg of protein.

All samples were reduced using 20 mM DTT in 50 mM TEAB at 50°C for 10 min, cooled to room temperature (RT) and held at RT for 10 minutes, then alkylated using 40 mM iodoacetamide in 50 mM TEAB at RT in the dark for 30 minutes. Samples were acidified with 12% phosphoric acid to obtain a final concentration of 1.2% phosphoric acid. S-Trap buffer consisting of 90% methanol in 100 mM TEAB at pH ∼7.1, was added and samples were loaded onto the S-Trap micro spin columns. The entire sample volume was spun through the S-Trap micro spin columns at 4,000xg and RT, binding the proteins to the micro spin columns. Subsequently, S-Trap micro spin columns were washed twice with S-Trap buffer at 4,000xg and RT and placed into clean elution tubes. Samples were incubated for 60 minutes at 47°C with sequencing grade trypsin (Promega, San Luis Obispo, CA) dissolved in 50 mM TEAB at a 1:25 (w/w) enzyme:protein ratio. Afterwards, trypsin solution was added again at the same ratio, and proteins were digested overnight at 37°C.

Peptides were sequentially eluted from S-Trap micro spin columns with 50 mM TEAB, 0.5% formic acid (FA) in water, and 50% acetonitrile (ACN) in 0.5% FA. After centrifugal evaporation, samples were resuspended in 0.2% FA in water and desalted with Zip Tips containing a C_18_ disk (MilliporeSigma, Burlington, MA). The desalted eluents were then subjected to an additional round of centrifugal evaporation and re-suspended in 0.2% FA in water at a final concentration of 1 µg/µL for fraction 1, 200 ng/µL for fractions 2, 3, and 4, and 1 µg/µL for Ex Vivo protein insolubilization assay samples. For fraction 1 and Ex Vivo protein insolubilization assay samples, four microliters of each sample was diluted with 2% ACN in 0.1% FA to obtain a concentration of 200 ng/µL. 0.5 µL of indexed Retention Time Standard (iRT, Biognosys, Schlieren, Switzerland) was added to each sample, thus bringing up the total final volume to 10 µL^75^.

### Mass spectrometric proteomics analysis

Reverse-phase HPLC-MS/MS analyses were performed on a Dionex UltiMate 3000 system coupled online to an Orbitrap Exploris 480 mass spectrometer (Thermo Fisher Scientific, Bremen, Germany). The solvent system consisted of 2% ACN, 0.1% FA in water (solvent A) and 80% ACN, 0.1% FA in ACN (solvent B). Digested peptides (400 ng for fractions 2, 3, and 4, and Ex Vivo protein insolubilization assay samples; 800 ng for fraction 1) were loaded onto an Acclaim PepMap 100 C_18_ trap column (0.1 x 20 mm, 5 µm particle size; Thermo Fisher Scientific) over 5 min at 5 µL/min with 100% solvent A. Peptides (400 ng for fractions 2, 3, and 4, and ex-vivo samples; 800 ng for fraction 1) were eluted on an Acclaim PepMap 100 C_18_ analytical column (75 µm x 50 cm, 3 µm particle size; Thermo Fisher Scientific) at 300 nL/min using the following gradient: linear from 2.5% to 24.5% of solvent B in 125 min, linear from 24.5% to 39.2% of solvent B in 40 min, up to 98% of solvent B in 1 min, and back to 2.5% of solvent B in 1 min. The column was re-equilibrated for 30 min with 2.5% of solvent B, and the total gradient length was 210 min. Each sample was acquired in data-independent acquisition (DIA) mode^58, 59, 76^. Full MS spectra were collected at 120,000 resolution (Automatic Gain Control (AGC) target: 3e6 ions, maximum injection time: 60 ms, 350-1,650 m/z), and MS2 spectra were collected at 30,000 resolution (AGC target: 3e6 ions, maximum injection time: Auto, Normalized Collision Energy (NCE): 30, fixed first mass 200 m/z). The isolation scheme consisted of 26 variable windows covering the precusor ion range of 350-1,650 m/z range with an overlap of 1 m/z between windows^76^.

### DIA data processing and statistical analysis

Insolublome DIA data was processed in Spectronaut (version 15.7.220308.50606) using the directDIA workflow. Data extraction parameters were set as dynamic and non-linear iRT calibration with precision iRT was selected. Data was searched against the *Mus musculus* reference proteome with 58,430 entries (UniProtKB-TrEMBL), accessed on 01/31/2018. Trypsin/P was set as the digestion enzyme and two missed cleavages were allowed. Cysteine carbamidomethylation was set as a fixed modification while methionine oxidation and protein N-terminus acetylation were set as dynamic modifications. Identification was performed using 1% precursor and protein q-value. Quantification was based on the peak areas of extracted ion chromatograms (XICs) of 3 – 6 MS2 fragment ions, specifically b- and y-ions, with local normalization and q-value sparse data filtering applied. In addition, iRT profiling was selected. Differential protein expression analysis comparing control to ketone conditions were performed using a paired t-test, and p-values were corrected for multiple testing, using the Storey method^77^. Specifically, group wise testing corrections were applied to obtain q-values. Protein groups with at least two unique peptides, q-value < 0.05, and absolute Log_2_(fold-change) > 0.58 are significantly altered.

### DDA library generation and DIA quantification for *ex vivo* protein insolubilization assay

A DDA spectral library was generated in Spectronaut (version 15) using BGS settings and the same *Mus* musculus database as stated above. Briefly, for the Pulsar search, trypsin/P was set as the digestion enzyme and 2 missed cleavages were allowed. Cysteine carbamidomethylation was set as a fixed modification, whereas methionine oxidation and protein N-terminus acetylation were variable modifications. Identifications were validated using 1% false discovery rate (FDR) at the peptide spectrum match (PSM), peptide and protein levels, and the most confident 3 – 6 fragments per peptide were kept. The spectral library contains 42,354 peptides and 3,862 protein groups. Identification was performed requiring a 1% q-value cutoff on the precursor ion and protein levels. Ex Vivo protein insolubilization assay DIA data was processed in Spectronaut (version 15) using the spectral library previously described just above from the acquired DDA acquisitions, and the same parameters as for the directDIA search. In addition, differential protein expression analysis comparing 1) 0 mM R-BHB to 1 R-BHB or 2) 0 mM R-BHB to 5 mM R-BHB were performed using the same parameters employed in directDIA search.

### Bioinformatics and proteomics visualization

Original datasheets were received from Buck Institute for Research on Aging Proteomics Core. Protein target UniProt AC/IDs, log_2_FC, and Q-Value (FDR) were imported into R Studio (Version 4.2.2, 2022_10_31, “Innocent and Trusting”) for downstream cluster and enrichment analyses.

Cluster analysis via partial least square-discriminant analysis (PLS-DA) of the proteomics data was performed using the package mixOmics^78^. Volcano plots were created using the package EnhancedVolcano^79^. Venn diagrams were created using the package ggvenn. Imported datasets were cleaned so only the primary UniProt AC/ID was used as a downstream identifier. AC/IDs were exported from R to UniProt for mapping to Gene Names and imported back to R for conversion to Entrez IDs using ClusterProfiler^80, 81^. ClusterProfiler was used for gene ontology (GO) biological process and Kyoto Encyclopedia of Genes and Genomes (KEGG) overrepresentation analysis (ORA). Background (denominator of GO and KEGG ORA) was whole mouse genome from package org.Mm.eg.db^82^.

GO and KEGG ORA results were visualized with dotplots using the package ggplot2^83^. KEGG ORA results were also used to manually curate BRITE hierarchical information for clustering visualization. Data were exported to GraphPad Prism for further visualization before final formatting in Biorender.

For the protein domain enrichment analysis, Interpro protein domain annotations were extracted from UniProtKB/Swiss-Prot (uniprot_sprot.dat, downloaded 2022_06_30) and mapped to each UniProt AC/ID using a custom Perl script (Perl v5.30.2). The frequency of domains among the entire set of mouse (species 10090) uniprot_sprot proteins was then used as the comparison for calculating fold-enrichment and binomal probability of the frequency with which each Interpro domain was found among each list of proteins (UniProt AC/IDs) in our proteomics data sets. Multiple-hypothesis correction is via Benjamini-Hochberg FDR^84^.

P Value calculated for one-tailed probability on Venn Diagram is as follows: 𝑧 = ((𝐾 − 𝑛𝑝) ± 0.5)/√𝑛𝑝𝑞, with K = 296 (actual targets found), n = 14635 (sum of detectable proteomes in all *ex vivo* and *in vivo* experiments), p = 0.2 (sum of shared targets between 1/0 and 5/0 R-βHB *ex vivo* groups and BH-BD targets, divided by n), and q = 0.8. We found that the z score = −54.4, and described the p value as 0.000001.

### Mass spectrometric proteomics data availability

Raw data and complete MS data sets have been uploaded to the Mass Spectrometry Interactive Virtual Environment (MassIVE) repository, developed by the Center for Computational Mass Spectrometry at the University of California San Diego, and can be downloaded using the following link: https://massive.ucsd.edu/ProteoSAFe/dataset.jsp?task=619d08c21b0f48e09fc3fabfed3c2ac7 (MassIVE ID number: MSV000091514; ProteomeXchange ID: PXD040985).

Enter the username and password in the upper right corner of the page: Username: MSV000091514_reviewer, Password: winter.

